# Identifying similar populations across independent single cell studies without data integration

**DOI:** 10.1101/2024.09.27.615367

**Authors:** Óscar González-Velasco, Malte Simon, Rüstem Yilmaz, Rosanna Parlato, Jochen Weishaupt, Charles D. Imbusch, Benedikt Brors

## Abstract

Supervised and unsupervised methods have emerged to address the complexity of single cell data analysis in the context of large pools of independent studies. Here, we present ClusterFoldSimilarity (CFS), a novel statistical method design to quantify the similarity between cell groups acroos any number of independent datasets, without the need for data correction or integration. By bypassing these processes, CFS avoids the introduction of artifacts and loss of information, offering a simple, efficient, and scalable solution. This method match groups of cells that exhibit conserved phenotypes across datasets, including different tissues and species, and in a multimodal scenario, including single-cell RNA-Seq, ATAC-Seq, single-cell proteomics, or, more broadly, data exhibiting differential abundance effects among groups of cells. Additionally, CFS performs feature selection, obtaining cross-dataset markers of the similar phenotypes observed, providing an inherent interpretability of relationships between cell populations. To showcase the effectiveness of our methodology we generated single-nuclei RNA-Seq data from the motor cortex and spinal cord of adult mice. By using CFS, we identified three distinct sub-populations of astrocytes conserved on both tissues. CFS includes various visualization methods for the interpretation of the similarity scores and similar cell populations.

## Introduction

Single-cell sequencing is a technology that captures the biomolecular status at cell resolution and measures multiple cellular markers in hundreds to millions of cells. Researchers can elucidate the genomic, epigenomic, and transcriptomic heterogeneity and variability in cellular populations and subpopulations (1).

Cell classification and labeling based on phenotypic characteristics is one of the key features of single-cell analysis, they are achieved by computationally grouping cells based on common gene expression patterns, leading to a compendium of known specific markers from previously discovered cell types, cell status (e.g., cell cycle markers), and developmental stages (2). Cell labeling in groups is a crucial task for determining the composition and heterogeneity that helps to understand tissue composition, microenvironment function, disease, and cell fate. The available number of large-scale single-cell datasets generated using a wide range of technologies is continuously increasing with the advent of public databases (3,4). However, wide data integration remains difficult, because of the inherent batch-specific systematic variations and high heterogeneity of single-cell data and because of data privacy regulations to aggregate human raw data. Bioinformatic removal of batch effects can lead to the loss of biological signals owing to a general assumption of identical biological conditions (5). Several specific tools for integrative analysis exist; however, each tool has its own advantages and limitations, without a clearly superior method (6,7). This is particularly relevant for obscured cellular populations, which are often underrepresented or highly specialized subcellular groups where robust markers have not been defined (8). In all these cases, the inclusion of additional data and integration can improve the characterization of understudied cell groups; however, recently proposed approaches and benchmarks show moderate accuracy in cell type labeling tasks on ground-truth datasets (9,10). Overall, integrative methods exhibit considerable performance variability and are highly susceptible to dataset peculiarities (Luecken et al. 2021). These issues have led to increasing attention and growing concerns regarding reproducibility between single-cell studies (12–15).

Here, we present ClusterFoldSimilarity, a new methodology that effectively analyzes the similarity between groups of cells from any number of different and heterogeneous datasets without the need for batch-effect removal or integration. Our method is based on the differences in abundance of the sequenced signal between cell groups, using a Bayesian inference approach and permutation analysis to obtain the fold-changes, and computation of similarity values based on the vector space of these differences in abundance. The underlying assumption is that the overall signal expression changes across different cell populations, subpopulations, or phenotypes in each individual dataset should be conserved in external studies with similar populations when using the same frame of reference as a comparison.

We show that our method is highly versatile and accurate and can be used to label single cell-data using reference datasets, including labeling cell types in scATAC-Seq data using scRNA-Seq or scATAC-Seq reference datasets as ground truth, with potential multimodal data analysis applications. Furthermore, we show how it can be used to compare groups of cells from different tissues and different organisms in cross-species studies, and to infer, to some extent, cell mixtures in heterogeneous clusters. Because our computational similarity method can handle any number of independent datasets, it will enable the analysis of large existing collections of single-cell studies, and it will contribute to solve the modern community efforts on building wide-scale single-cell atlases across tissues, organs and organisms (16–18)

Finally, we exemplify how our methodology can be used to study small cell subpopulations across tissues by sequencing single-nucleus RNA-Seq from the spinal cord and motor cortex of adult mice and analyzing astrocytes as a disease-relevant glial cell subpopulation.

## Results

### Cluster Fold Similarity Overview

ClusterFoldSimilarity computes scores for each pair of cell groups based on differences in abundance, these values are centered around 0 (negative scores mean dissimilarity, while positive values are similar phenotypical states based on abundance differences). A summary of the workflow for computing the similarity metric is shown in Fig. 1. The tool expects two or more single-cell datasets, each with their raw feature count matrix, and defined groups of interest at the cell level. The identity of these groups can be a phenotypic characteristic (e.g., cell type, cell cycle, or time series) or an algorithmically obtained label (e.g., clustering techniques). In the first step, differences in abundance from raw expression between the groups in each dataset are estimated using a Bayesian approach. These fold-changes are normalized and used to compute a similarity metric between groups from independent single-cell assays. Additionally, the contribution of each feature to the similarity score is used to understand its importance as a marker of these specific groups of cells, acting as a feature importance discovery tool. To understand the complex interaction between the groups and their similarities, CFS builds a directed graph using the igraph framework, in which edges represent the direction (arrow) and intensity (length) of similarities from cluster source to cluster target. An additional community algorithm can be applied to find communities of clusters from the similarity graph, helping to identify the most similar cell communities.

**Figure 1:**
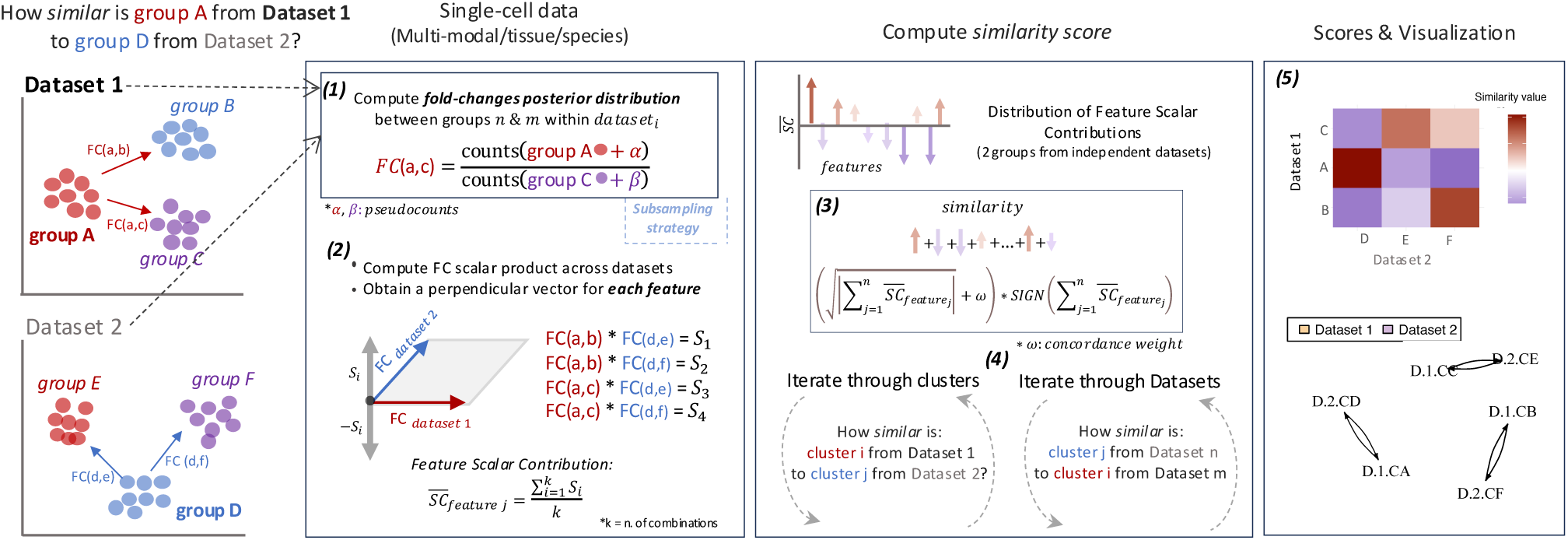
Diagram of ClusterFoldSimilarity for the comparison of two independent datasets. The datasets are depicted by dimensionality reduction (PCA, UMAP, t-SNE, VAE, PATHE or others) for visualization purposes only.

### Comparison with other popular integrative methods

ClusterFoldSimilarity can process a total of 750k cells, divided into three datasets with 20 clusters each (k=6819 combinations) in three minutes using seven cores (Fig. 2a). To assess the performance of CFS against other popular methods, we first selected two public annotated datasets and computed the accuracy of predicting the cell type by cluster similarities. The first scenario is composed of two scRNA-Seq pancreatic tissue datasets, from mouse and human samples (19). The second scenario uses an assay of scRNA-seq that aims to study mouse gastrulation across entire embryos on E8.5 timepoint (20). These datasets were used as ground-truth reference and unlabeled query iteratively, with cell groups defined based on their known cell-type annotation by the original authors. For the pancreatic scRNA-Seq, apart from the organism and sample batch effect (human samples n=4, mice individuals n=2), these datasets have a large difference in the number of available cells (human dataset n=7561 cells, mice dataset n=1861 cells), which allowed us to benchmark the sample size impact on the performance (Fig. 2b), and show that on average CFS is not significantly impacted by the varying size of the datasets. We performed an extensive benchmark comparison with other integrative and batch correction algorithms: MultiMap; naive PCA with no batch correction, PCA corrected using MNN (mutual nearest neighbors), and PCA corrected using Harmony; Seurat CCA (canonical correlation analysis); StabMap with no batch correction, StabMap corrected using MNN. We iterated across all datasets, using each as the reference and the query, and with varying number of available features (n=25, 50, 100, 250, 500, 1000, 1500) following a similar approach as depicted by Ghazanfar et. al. (10). In our case, for each reference-query pair we select from the top 2000 high variable features n random genes more than once (producing several predictions for the same reference-query n features pair), to assess the accuracy for different sets of genes. Results show high accuracy of CFS at cell type mapping task, especially when low number of shared features are available (Fig. 2c,d), case in which CFS has a better performance, highlighting that it can be applied in scenarios where the number common shared features is low, like multi-organism or multi-modal (e.g. single-cell proteomics with extremely small number of surface markers screened).

**Figure 2.**
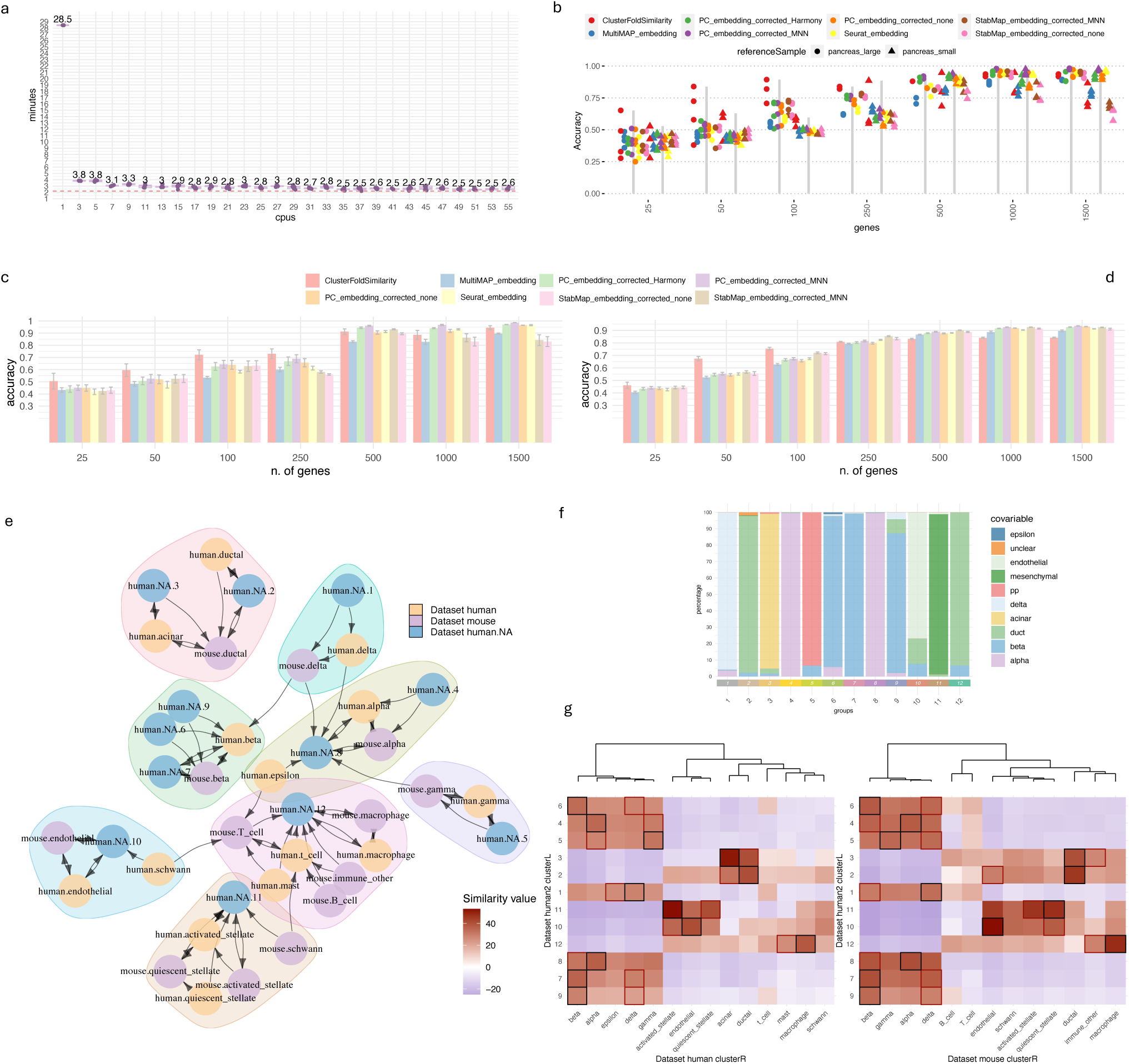
a. Benchmark of computing times of CFS using different numbers of CPUs. The artificial dataset is composed of 750k cells, divided into three datasets with 20 clusters each (k=6819 combinations). b. Plot showing cell type classification accuracy benchmark comparison against integrative methods (MultiMap; naive PCA, PCA MNN corrected, PCA Harmony corrected, Seurat CCA, StabMap, StabMap MNN corrected) using two pancreatic scRNA-Seq datasets with varying sizes. c. Barplot displaying mean cell type classification accuracy results of the Benchmark comparison against other integrative methods using two pancreatic scRNA-Seq datasets, each gene subset includes several permutations per gene sampling. Error bars represent the standard error of the mean. d. Barplot showing mean accuracy results of cell type classification benchmark comparison against integrative methods using a mice gastrulation scRNA-Seq datasets (n=4 mouse) on E8. Error bars represent the SEM. e. Directed graph of clusters by top similarity; each node corresponds to a cluster/cell type group from one dataset (human and mice). Length of edges corresponds with the magnitude of the similarity and arrows depict the direction of the similarity test. **f**. Proportion of cell types per cluster using the authors annotation (Muraro et. al,) from human pancreatic single cell dataset used as an unannotated target (human.NA). g. Heatmap showing all computed similarity scores (all possible combinations of groups) between the datasets.

In general, results suggest that the performance and accuracy of integrative single-cell methods are dataset-dependent, as an example StabMap showed a varying performance, being one of the topmost accurate in the gastrulation dataset (Fig 2d), but one of the worst performing in the pancreatic multi-species dataset (Fig 2c). This again highlights that a careful consideration of the selected method should be done for each independent single-cell data analysis, and many factors must be considered beforehand, like the number of features available, size of the datasets, degree of tissue/organism/technical batch effects, and others.

### ClusterFoldSimilarity effectively compares groups of mice and human cells

We used one mouse and two publicly available human single-cell RNA-Seq datasets from pancreatic tissue, each with annotated cell-type labels, to evaluate the accuracy of the similarity metric. The first human dataset (21) was subjected to unsupervised clustering to interrogate the cell-type composition, retaining the annotated cell types for later comparison. The other two datasets, from mouse and human pancreatic samples (19), were used as ground-truth references, with cell groups defined based on their known cell-type annotation by the original authors.

To assess the pair matches between the clusters in the unannotated human dataset and the cell type groups in the remaining studies, we computed and selected the top similarity scores for each of the 11 unannotated clusters (Supplementary table 1). This approach allowed us to confidently establish matches between clusters and annotated cell type groups (Fig. 2e,f,g). The results show that 10 out of 11 clusters exhibited perfect matches to the known annotations of the clusters from the original publication and both reference datasets.

Notably, the only exception (cluster 3), corresponding to acinar cells, displayed the highest similarity with the acinar cell group from the second human dataset and the ductal cell group from the mouse dataset. This could be expected, as acinar cells were absent in the annotation of the utilized mouse dataset. Notably, our similarity metric successfully indicated the closest related cell type by lineage: ductal cells. Furthermore, we observed a consistent association for cluster 11, which represents mesenchymal cells (a cell type that is not present in both reference datasets), with stellate cells, which were missing in the clustered human dataset, but present in both mouse and second human reference datasets. This association has been previously demonstrated in the literature (22).

Our results demonstrated the capability of our algorithmic approach to successfully match comparable phenotypic cell groups, even when working with different organisms. Notably, using more than one dataset as a reference helped to clear and define the broad relationships between cell populations (Fig. 2e); we believe that ClusterFoldSimilarity will benefit for the numerous public cell atlases available. However, it is important to acknowledge the potential limitation of our metric, which operates at the cell group level rather than at the individual cell level, which may lead to some loss of resolution in the annotation of heterogeneous clusters. However, we believe that this issue can be mitigated to some extent by increasing the granularity in the clustering analysis or by using existing meta-cell analysis algorithms that capture small cell groups with the same technical variability and thus maximize data resolution (23–25).

### Correlation between cell mixture proportion and similarity score

To investigate the effect of different cell mixtures within clusters on our similarity metric, we generated an artificial single-cell RNA-Seq dataset using the scDesign2 algorithm (26). The data were constructed based on a human pancreatic dataset (19) with known cell type labels. We adjusted the artificially recreated count matrix to ensure an identical number of cells per cell type (n=1200), thereby minimizing the cluster size effect and fostering the signal of small-cell populations (Supplementary Fig. 1a).

Using CFS, we computed the similarity scores between clusters from the original human pancreatic dataset, which included clusters with cell type mixtures, and the artificially recreated and cell type balanced dataset. Interestingly, the similarity scores demonstrated the ability to effectively capture cell mixture proportions at some extent (Supplementary Fig. 1b). For example, in the original dataset, Cluster 10 comprised up to nine different cell types, with marginal cell types not typically found in the pancreas. The similarity scores mirrored the cell type proportions in descending order: endothelial cells, macrophages, mast cells, Schwann cells, T-cells (Supplementary Fig. 1b). Only stellate cells exhibited a higher similarity value than anticipated, although they were still lower than those of the top two most abundant cell types. Similar trends were observed in Cluster 4 of the original dataset, composed of five different cell types. The similarity values again correlated with the percentage of cells from each cell type in the following descending order: activated stellate cells, quiescent stellate cells, endothelial cells, acinar cells, and Schwann cells (Supplementary Fig. 1b,c,d).

Overall, our results demonstrate how CFS is able to capture -at some extent-the complex mixture of cell populations, although we do not believe that it will be suitable for deconvolutional single-cell task, we think it is an important point to take into consideration when analyzing and interpreting CFS results, and we believe similarity values exhibit great potential for downstream analysis, and can be integrated in combination with additional bioinformatics methods.

*Supplementary Figure 1. **a** UMAP dimensionality reduction of human pancreas Dataset 1 from Baron et. al (top) and simulated dataset (below) showing the cluster labels (left) and annotated cell type label (right). b Barplots showing the number of cells (blue) versus the similarity value for the cell type (yellow). Left: results for Cluster 10, right: results for Cluster 4. c Heatmap showing all the computed similarity scores between datasets. **d** Directed graph of clusters by similarity, where each node corresponds with a cluster/cell type group from one dataset (dataset original: dataset from Baron et. al, simulated: dataset simulated from the original using scDesign2)*.

### Expanding to multimodal assays: Single-Cell proteomics, sorted bulk RNA-Seq and single-cell RNA-Seq

To test whether cell populations from additional multimodalities can be linked using our technique, we analyzed 51 samples of synovial tissue biopsies from rheumatoid arthritis (RA) and osteoarthritis (OA) from sorted B cells, T cells, monocytes, and stromal fibroblast populations (27). The multimodal study included sorted bulk RNA-seq from all patients (n=54 patients), single-cell RNA expression (n=21), and single-cell protein expression from a 34-marker mass cytometry panel (Cytof, n=26).

We processed the three datasets independently, obtaining 13 clusters for the single cell RNA-Seq assay to be used as targets of the annotation by similarity, and used the cell type annotations from the original authors as ground truth labels for both bulk RNA-Seq and Cytof. Annotations for the computed single cell RNA-Seq clusters were retrieved and compared with the unannotated cluster for later comparison (fibroblast clusters 0, 4, 6; T cell clusters 1, 5, 8, 9, 12; B cells clusters: 2, 7, 11; monocyte clusters: 3, 10). The data processing differs considerably from the original study, as an integration strategy is used by the authors, which might negatively impact the clustering analysis especially in the case of cell subtypes.

The similarity metric was computed using 33 common features across all datasets (proteins in the case of mass spectrometry data and genes coding for those proteins in the case of RNA-Seq assays). For all the cell type classes present in the assays (B cells, T cells, monocytes, and fibroblasts) clusters of cells with the same pre-annotation were unambiguously mapped together using our similarity metric (Supplementary table 1) for all three data modalities, including the unannotated clusters (Supplementary Fig. 2a,b). Only the small cluster CD4+HLA-DR+ from the Cytof data, composed of T cells, was linked per top similarity to cluster 3 from single cell RNA-Seq, composed of monocytes. Although the same cluster 3 (monocytes) from single cell RNA-Seq assay showed as its top similarity the Cytof cell group CD11c+ CD38− CD64+ (Supplementary Fig. 2a), which is composed solely of monocytes. This shows that using more than two datasets can improve the similarity matching and thus the understanding of complex interactions between groups. Notably, even with an extremely small number of common features available (n=33), our approach demonstrates to be robust and accurate, showing again the flexibility and adaptability of the methodology.

*Supplementary Figure 2. a. Directed graph of top similarity values between clusters and cell groups from single cell RNA-seq, sorted bulk RNA-seq and Cytof samples from rheumatoid arthritis (RA) and osteoarthritis (OA) from sorted B cells, T cells, monocytes, and stromal fibroblast populations (single cell RNA-Seq fibroblast clusters 0, 4, 6; T cell clusters 1, 5, 8, 9, 12; B cells clusters: 2, 7, 11; monocyte clusters: 3, 10) b. Heatmap of all similarity values from single cell RNA-Seq clusters and bulk RNA-Seq and Cytof samples. c. Heatmap showing all computed similarity scores between the single cell ATAC-Seq gene activity scores and the RNA-Seq annotated dataset. Black squares highlight the top value of similarity, dark red squares the top second similarity value. d. Selected markers by ClusterFoldSimilarity for clusters of the ATAC-Seq data*.

### Similarity matches population across single-cell RNA-seq and ATAC-seq data

We applied our novel methodology to multimodal single-cell data comprising RNA-seq and ATAC-seq from two PBMC datasets. As a reference, we used a high-quality CITE-seq atlas of the circulating human immune system, comprising 160k cells with 26 annotated cell types (28). Additionally, we processed one non-annotated public 10x ATAC-Seq dataset, obtaining 18 clusters that were manually labeled by cell type using canonical markers, including fine annotations for specialized subtypes, with a total of 15 different phenotype characterizations. We then computed the similarity metric from the ATAC-Seq clusters against the reference ground-truth CITE-seq annotated data. For ATAC-Seq, the gene activity matrix was computed from peak calling using Signac, easing the mapping between features from different modalities and datasets.

Overall, similarity scores were able to link clusters from ATAC-Seq data to reference cell populations in RNA-Seq data with identical phenotypes (Supplementary Table 1), including small populations such as plasmacytoid Dendritic Cells (pDCs) and myeloid Dendritic Cells (mDCs), and differentiate between specialized subgroups of CD8 and CD4 of T-memory VS T-naïve (Supplementary Fig. 2c). Results showed that out of 17 comparable cell populations, 14 obtained top similarity values with the corresponding and matching cell types. The second and third most similar groups were the corresponding types from those that did not map with the most similar group. For instance, non-classical monocytes top similar group in the reference was classical monocytes, followed closely as second most similar by the non-classical monocytes. Interestingly, one cluster of CD8 T-memory cells was found to be most similar to Natural Killer cells, and KLRD1 selected by our tool (Supplementary Fig. 2d), the marker gene contributing the most to the score. KLRD1 is used as an NK marker and is known to be expressed in CD8 cytotoxic cells (29–32). The top markers selected by feature selection ability of CFS were in general highly specific (Supplementary Fig. 2d).

In the case of some highly specialized manually annotated groups that were not present in the reference dataset, cell types engaging in those specific phenotypes were found to be the most similar (e.g., the top similar group for the ATAC-Seq manually annotated T helper cell clusters was CD4 naïve from the reference single-cell RNA-Seq, Supplementary Fig. 2c).

These results show that our method is a powerful tool not only for matching clusters or finding markers of interest but can also be used in conjunction with high-quality reference datasets.

### Astrocyte analysis on mouse motor cortex and spinal cord

To further demonstrate the versatility of our metric, we sequenced and analyzed single-nuclei RNA-Seq from the motor cortex and spinal cord of three wild-type 12 months old mice, for a total of six samples (Fig. 3a), with the focus of identifying and characterizing astrocytes in different parts of the motor-neuronal system. Astrocytes are glial cells in the central nervous system with a wide diversity of roles and physiological functions, including development and maturation of synapses, neurotransmitter homeostasis, glycogen storage, blood brain barrier (BBB) maintenance, and clearance of protein aggregates (33–36). They have become of high interest as they contribute to several neurological diseases, such as the motor disease amyotrophic lateral sclerosis (ALS) (37–39). After processing and filtering for quality control, a total of 39493 cell-nuclei were selected for downstream analysis (mean of nuclei per mouse=6582, sd=1107). Astrocytic cell populations were selected using respective canonical markers (Fig. 3c), for a total of 3056 astrocytes (mean per mouse=509 astrocytes, sd=167).

**Figure 3.**
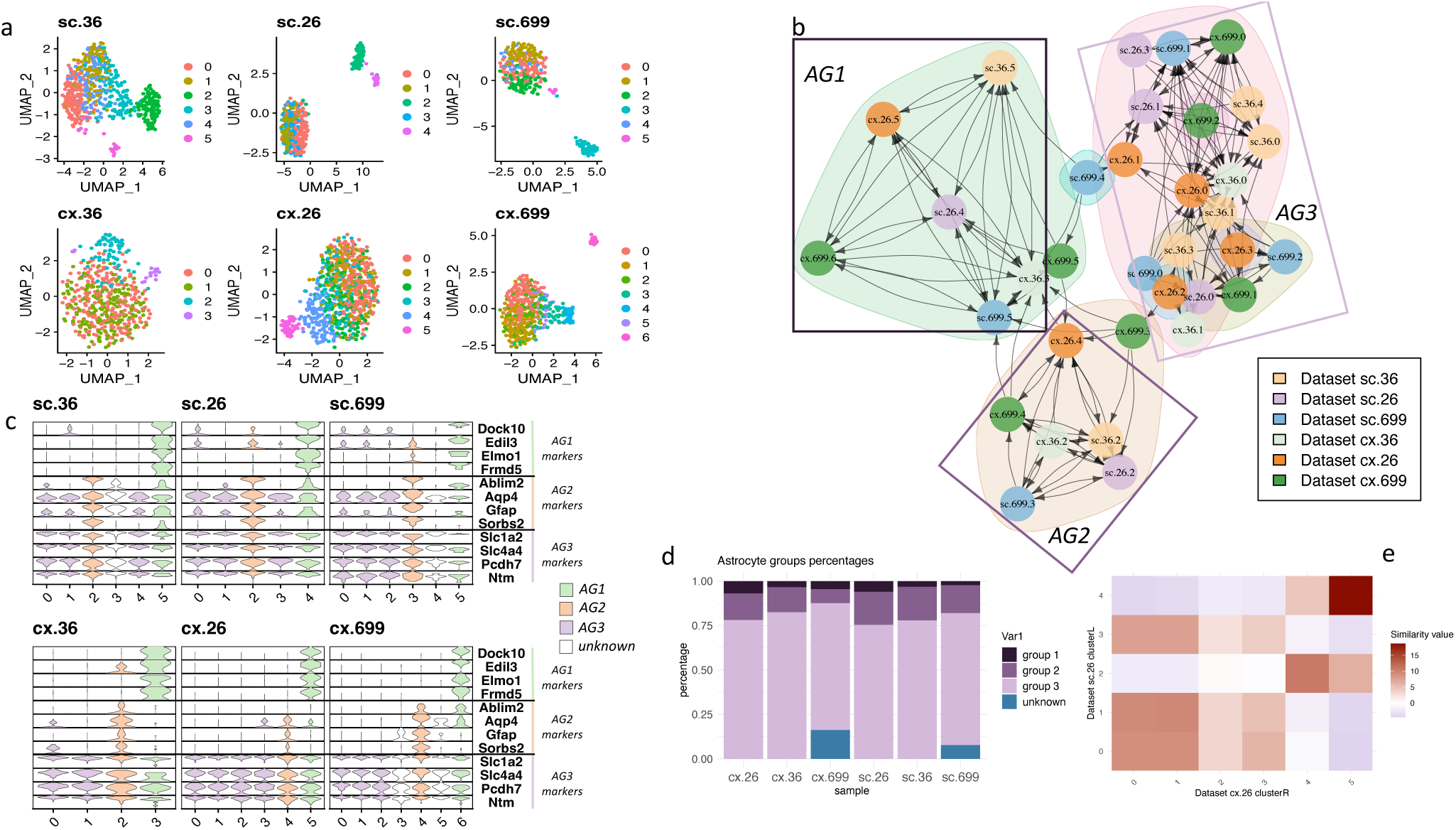
Integration of astrocyte cluster subpopulation using our similarity metric. a UMAP dimensionality reduction of datasets (N=6) sc:spinal cord, cx:cortex. b Directed graph of clusters by similarity, each node corresponds to a cluster from one dataset, edges correspond to the similarity value and the direction of the similarity, shorter edge lengths translate to larger similarity values. c Violin plots of astrocyte gene markers selected by CFS as contributing the most to the similarity value for the specified AG1, AG2 and AG3 subpopulations. d Percentage of cells corresponding to each astrocyte group per sample. e Heatmap plot of all the similarity values between spinal cord and cortex from the same mouse (id:26).

*Supplementary Figure 3. Integrated single-nuclei RNA-Seq data from spinal cord and motor cortex from the three mice. a. UMAP plot showing all cells by mice id. b. UMAP plot showing all cells by tissue id. c. UMAP plot showing astrocyte groups detected by ClusterFoldSimilarity in the context of all cells. d. UMAP plot showing astrocytes by mice id. e. UMAP plot showing astrocytes by tissue id. f. UMAP plot showing astrocytes sub-populations detected by ClusterFoldSimilarity*.

An initial integration of the expression data revealed, as expected, major differences between spinal cord and motor cortex (Supplementary Fig. 3b,e). To overcome the tissue and intra-specimen variability (Supplementary Fig. 3a,b,d,e) and identify conserved astrocyte populations, we used our similarity score analysis to match clusters between mice. For this, each sample was independently processed (Fig. 3a) and clustered (see Materials and Methods). The processed samples were then fed into our similarity score algorithmic approach, using all available genes, so that novel markers outside the top variable genes could be detected by CFS feature selection ability.

Exploratory analysis of common markers like Gfap and Aqp4 showed high variability (Fig. 3c), revealing the emerging heterogeneity and diverse functionality of astrocytes (40,41). Glial fibrillary acidic protein (Gfap), while showing overall expression in spinal cord, was cluster-specific in motor cortex, showing the limitations of this marker as previously described (42,43). In the same direction, Aquaporin 4 (Aqp4), highly expressed in astrocytes and responsible for water exchange across the brain, showed consistent expression in spinal cord astrocyte populations, but not in the motor cortex. The enzyme aldehyde dehydrogenase 1 family member L1 (Aldh1l1), regulating folate metabolism and known to be strongly expressed in astrocytes, showed overall inconsistent expression across tissues and clusters.

### Similarity score identifies conserved astrocyte populations across motor cortex and spinal cord

Using the similarity scores from the clusters of each sample and the graph community analysis using leiden algorithm, we defined 3 distinct subpopulations of astrocytes based on differences in transcript abundances (Fig. 3b-e). Isolated clusters with few or no inward arrows on the similarity graph were excluded (cx.699 clusters 3 and 5, sc.699 cluster 4, sc.36 cluster 3). These subpopulations were observed in both motor cortex and spinal cord. Top features selected by CFS as the ones contributing the most to the observed similarities were highly specific in most of the cases (Fig. 3c) and in they were observed in the significant DEG found on the contrast analysis withing each group (see next section). Astrocyte Group 3 (AG3) was the largest and most heterogeneous, comprising three additional sub-groups showcasing similar characteristics but different expression patterns. In contrast, AG1 was the smallest in terms of number of cells, yet exhibiting high homogeneity, and showing highly specific markers found by our similarity method. AG2 resembled GFAP+ physiologically active population, showing upregulated genes that are shared with pathogenic reactive populations (44).

To better understand the main differences between these three AG populations, we performed a differential expression analysis between the groups discovered by CFS pulling all the data together and adding as model covariates for the DE test the mouse id and sequencing date. A justification for this is that the marker selection capabilities of CFS are complementary to differential expression analysis, but a more refined test is needed to uncover the significant differences once the main groups have been identified, since features contributing to the similarity between clusters can be relevant for several cluster pairs and thus not necessarily differentially expressed between main groups.

### A small astrocyte population enriched in neuronal and CNS developmental markers

In total, we found 330 differentially upregulated genes (p. value < 0.005, see materials and methods statistical significance) in Astrocyte Group 1 (AG1) in contrast with AG2 and AG3 (Table 1), both in spinal cord and motor cortex tissues (differential expression analysis was done using the merged data with the tissue, sequencing date, and the mouse id as covariates). A large quantity of these upregulated genes were linked to neurogenesis and nervous system development functions (Fig. 4a,b). Ontology analysis revealed a set of major biological terms significantly associated with neurogenesis, nervous system development, neuron projection, neuron differentiation, and cell projection organization (Fig. 4b). Relevant actors in this upregulated list include a set of transcription factors (TF) uniquely expressed in AG1. Sox10 involved in embryonic development and cell fate, has been described to play an important role in neural crest and peripheral nervous system development (45), as well as activating additional transcription factors (46,47). Zfp536 with roles in regulation of neuron differentiation and negative regulation of retinoic acid receptor signaling pathway (48,49). Zeb2 (Sip1, Zfhx1b), a TF that acts both as a DNA-binding transcriptional repressor and activator, is known to be involved in neuronal development (50) and in the differentiation between cortical and striatal interneurons (51); Zeb2 shows expression across all groups in our study, being significantly higher in AG1.

**Figure 4.**
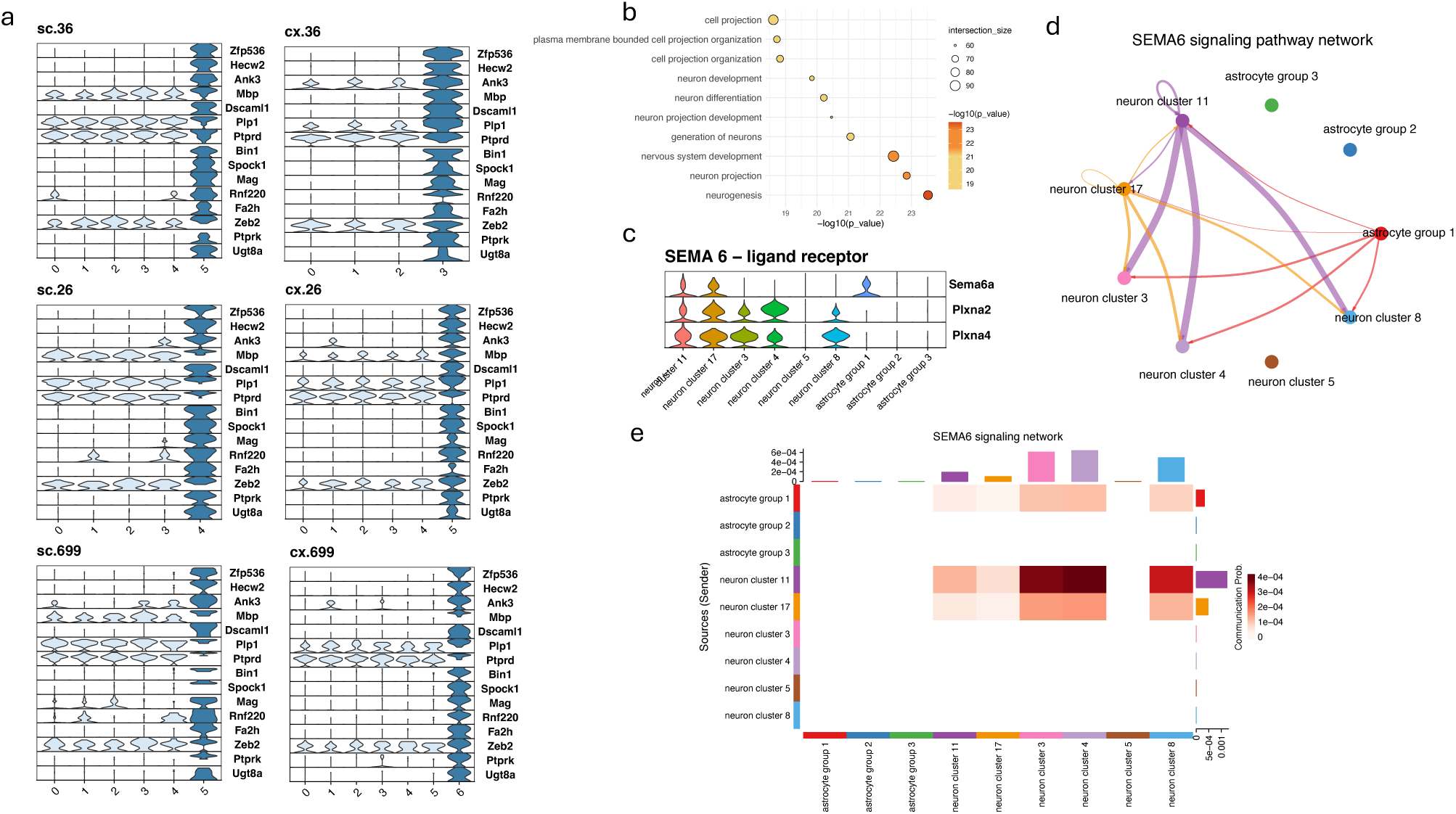
Detailed analysis of Astrocyte Group 1. **a.** Gene markers related with neurogenesis and neuronal development. Dark blue violin plots correspond to cell clusters of AG1 in individual samples (cx: cortex, sc:spinal cord). **b.** Gene ontology enrichment analysis results of the significantly up-regulated markers in AG1. **c.** SEMA6 signaling pathway ligand-receptor components RNA expression across neuronal clusters and astrocyte groups. **d.** SEMA6 cell-cell communication network analysis between neuronal clusters and astrocyte groups. The only astrocyte group actively communicating is AG1. e. Heatmap of the SEMA6 signaling pathway and the role in the communication of the different neuronal and astrocyte groups.

**Table 1.**
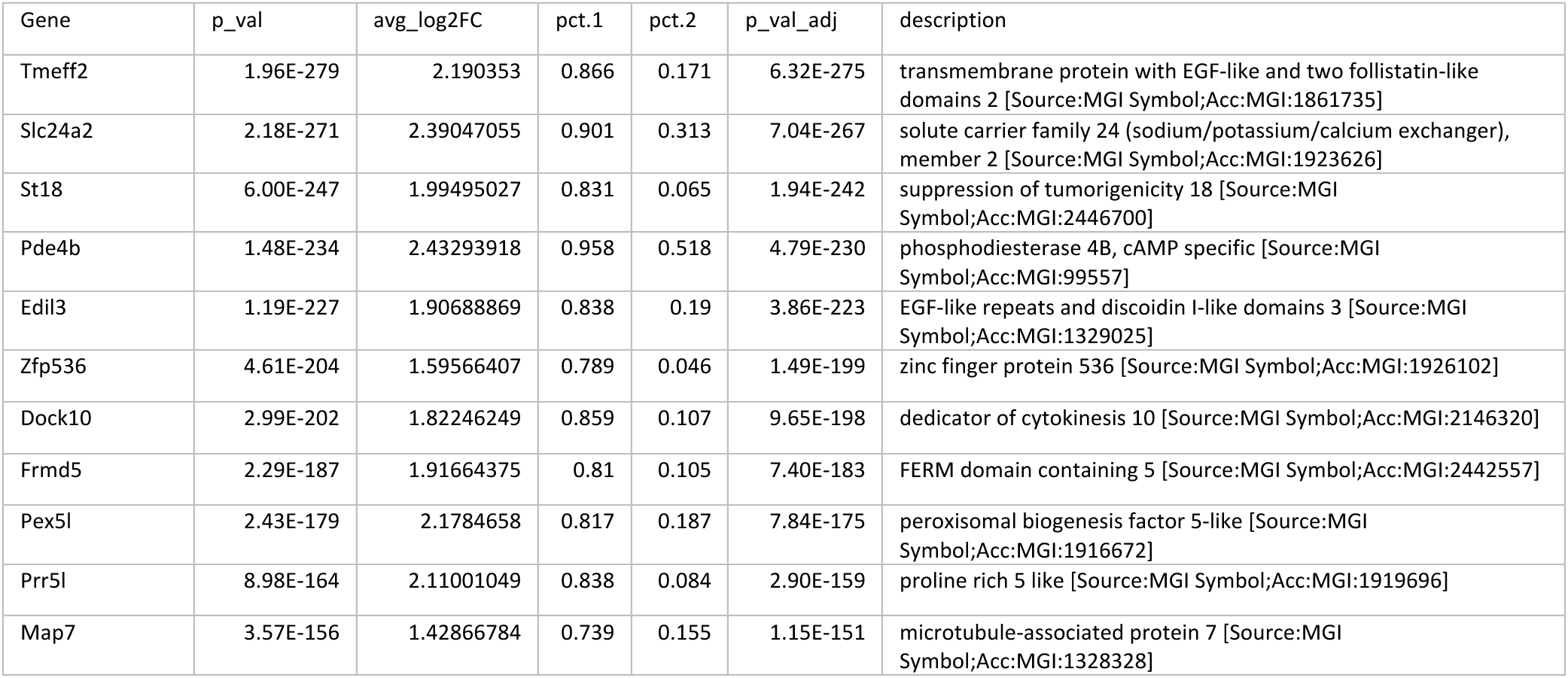
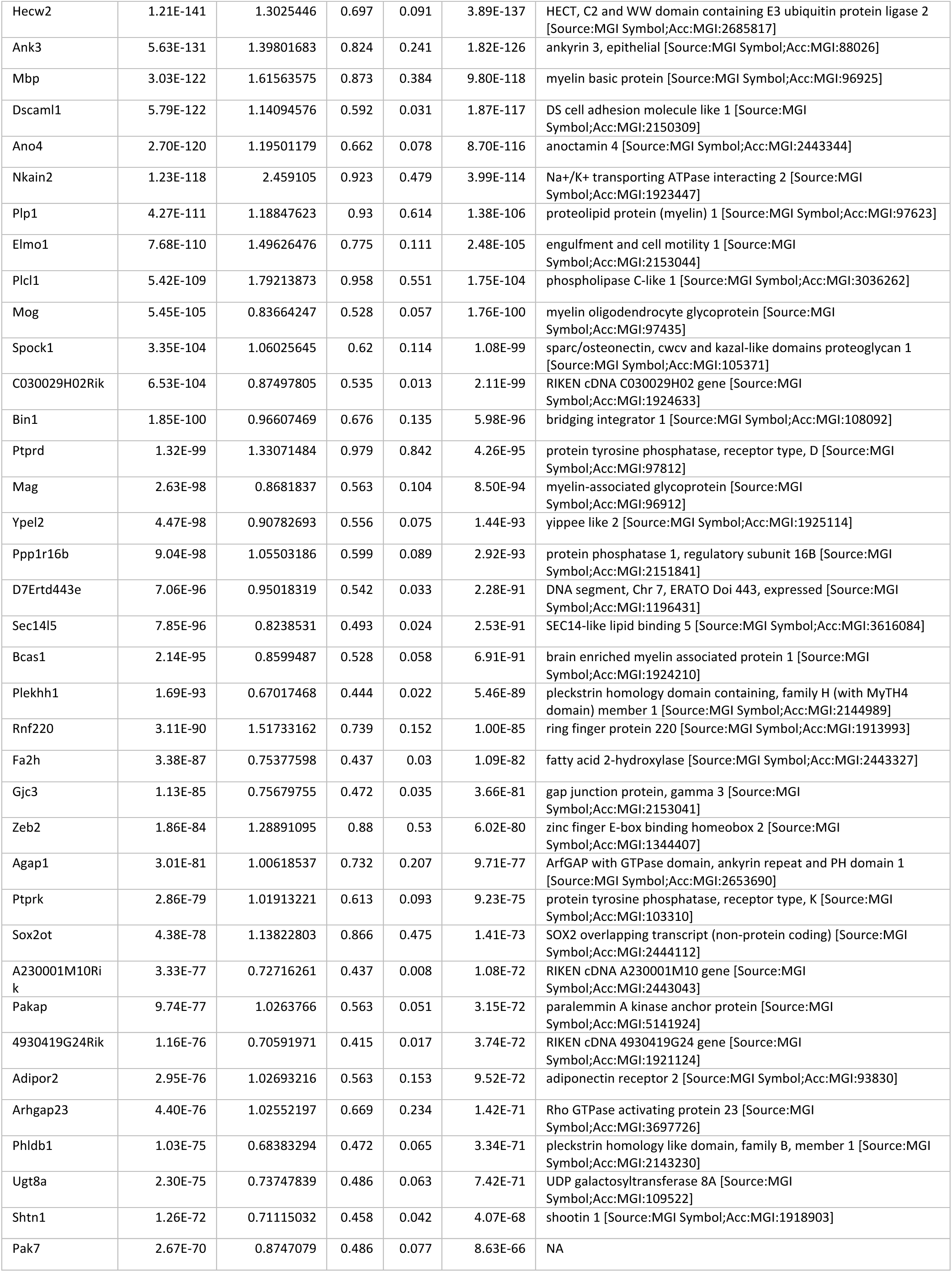
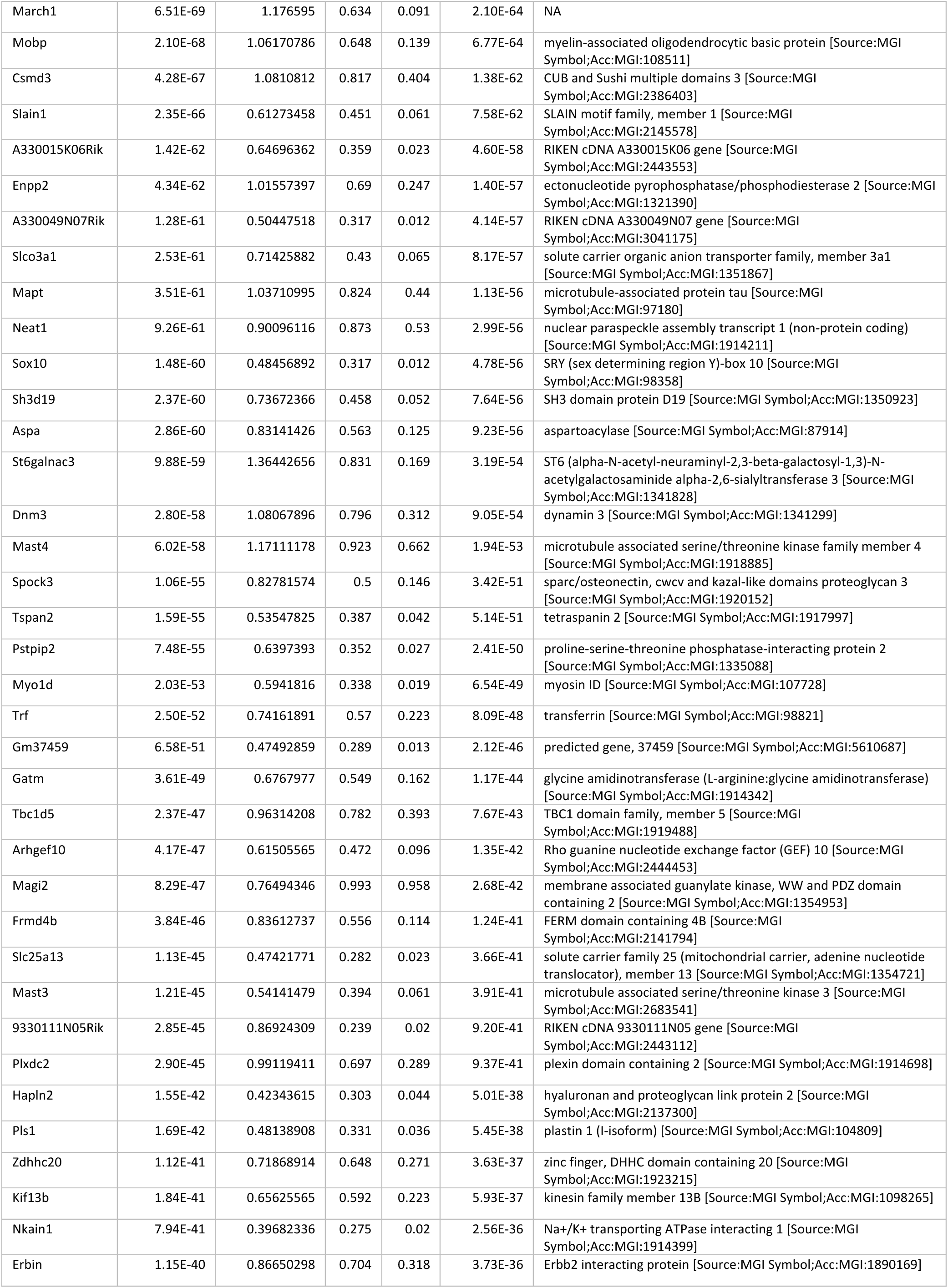
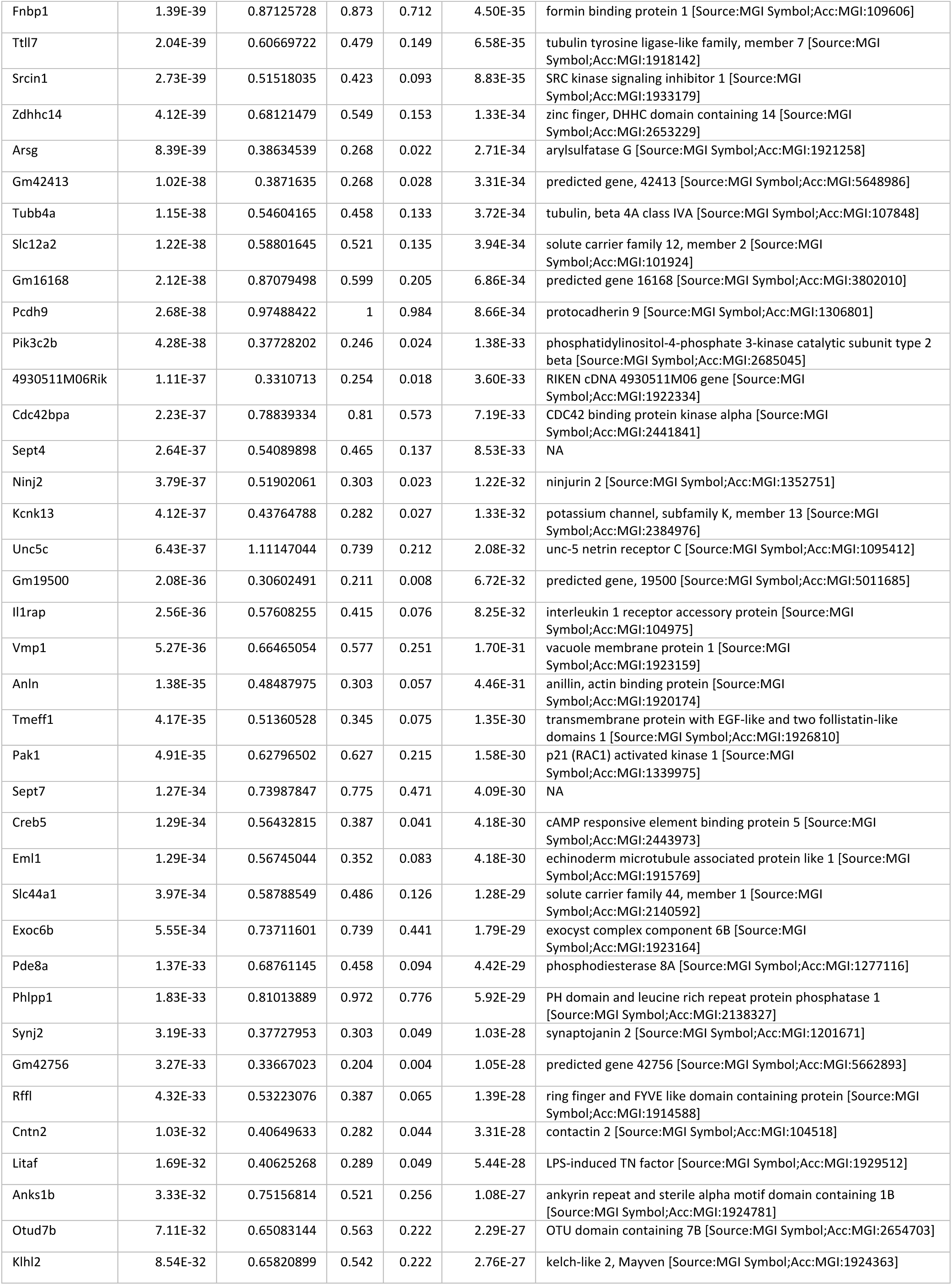
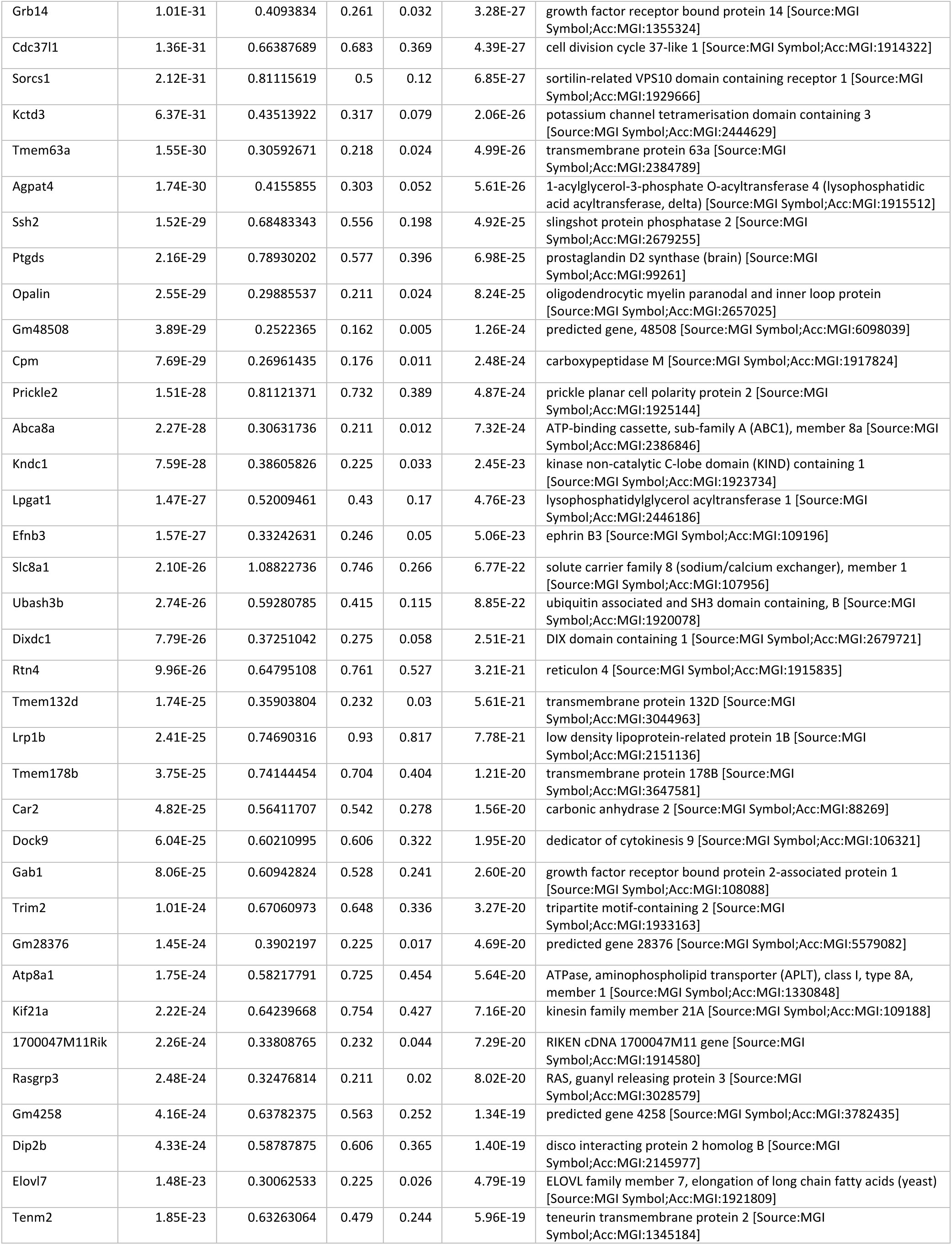
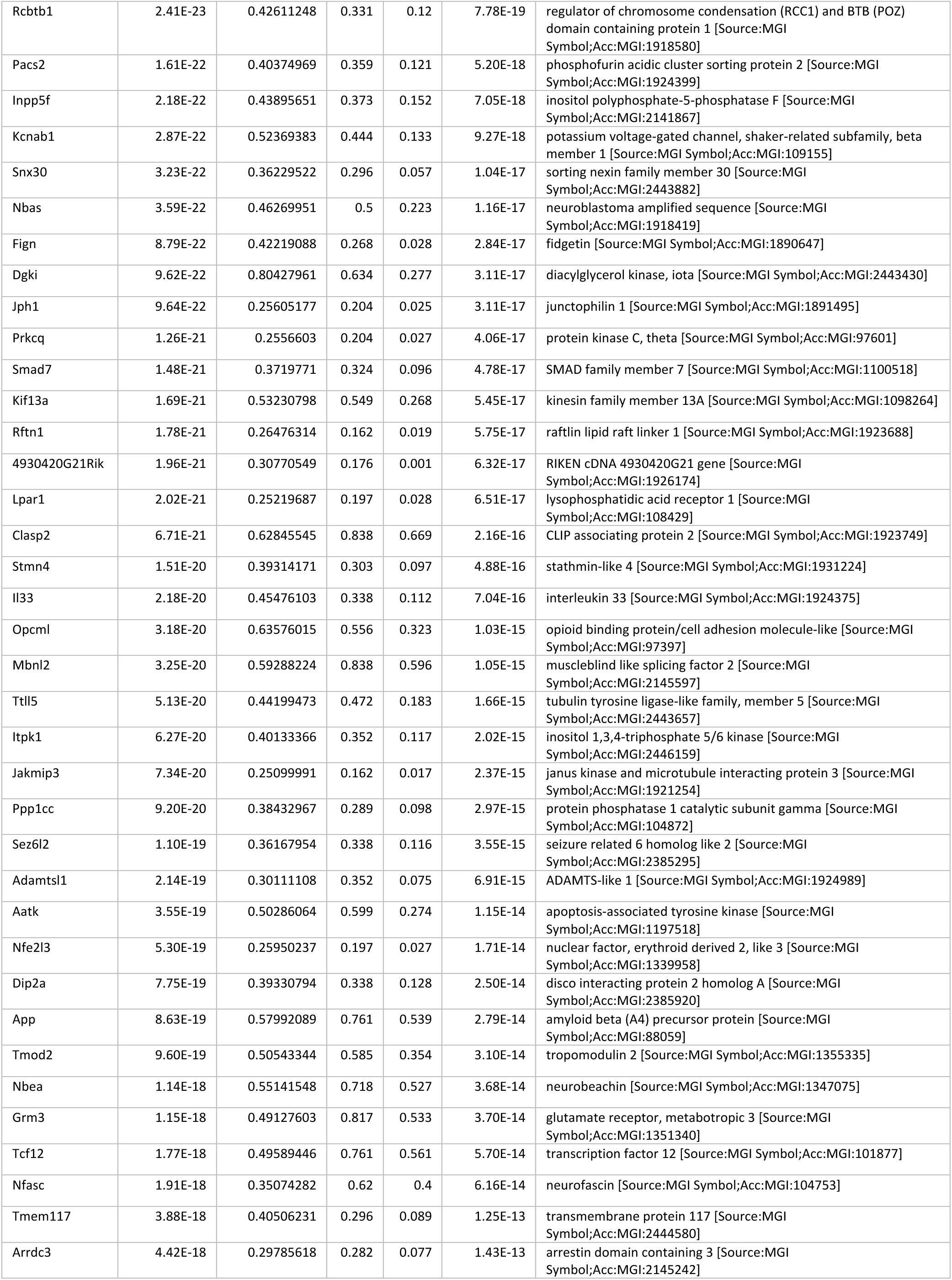
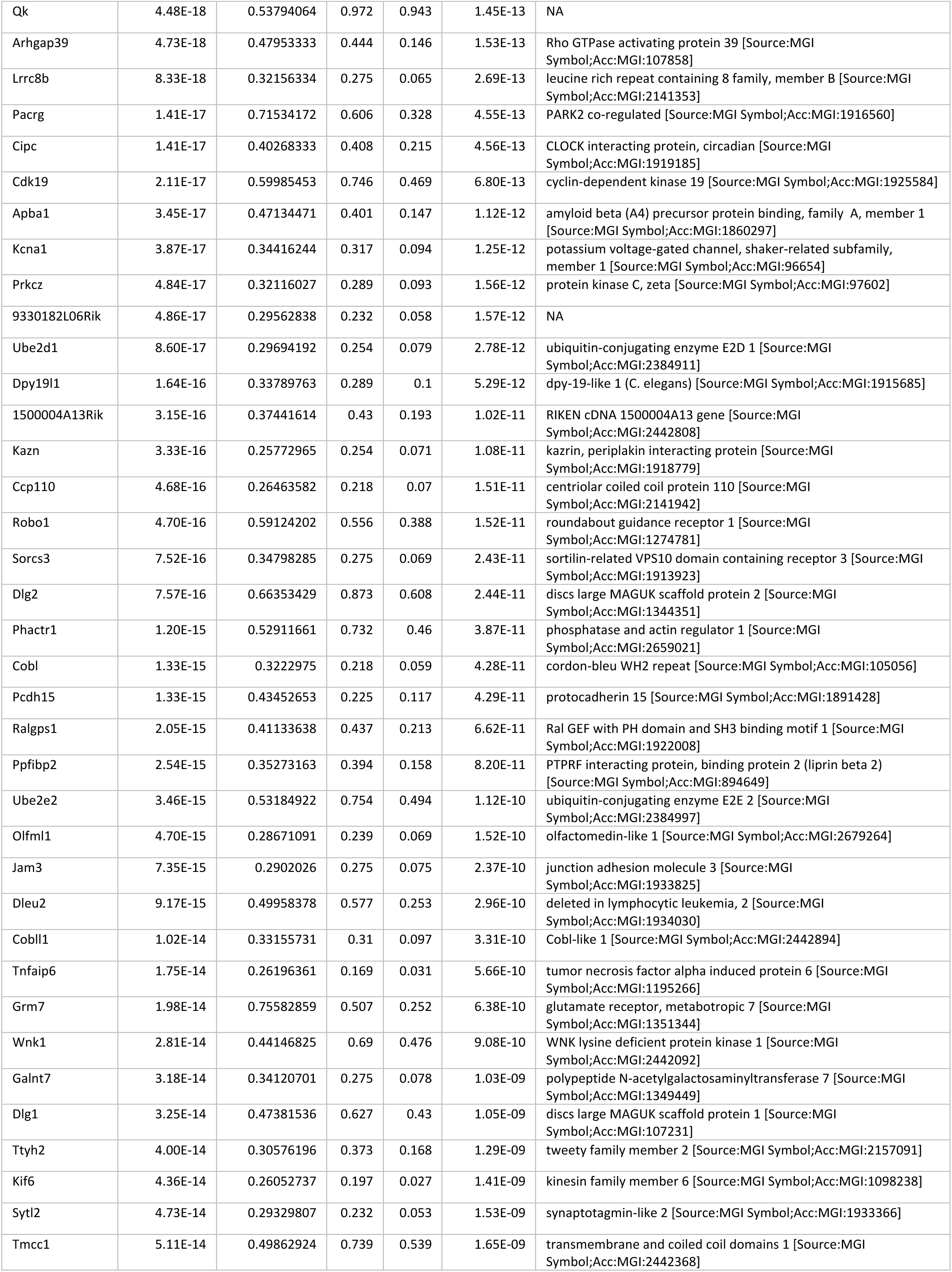
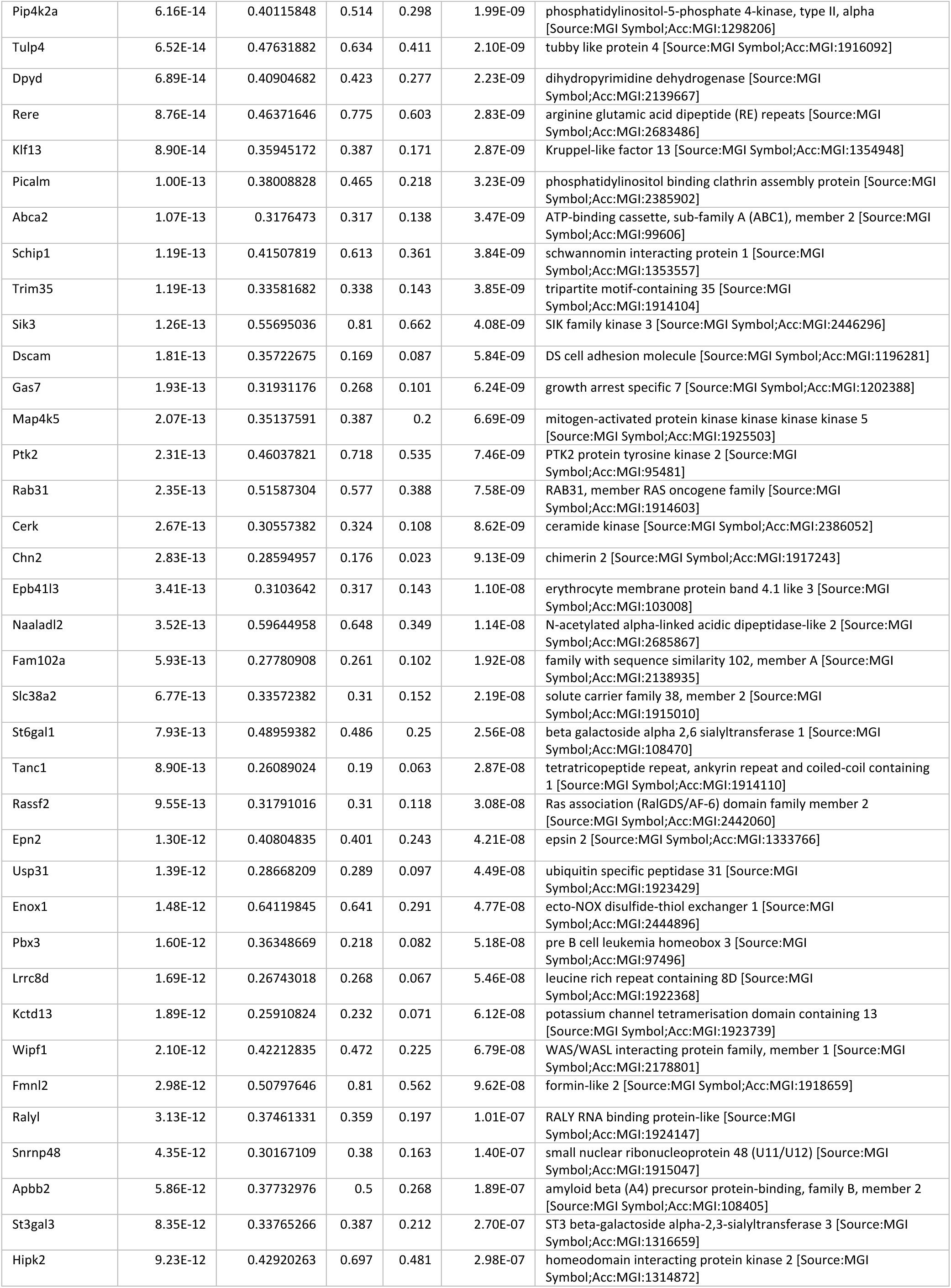
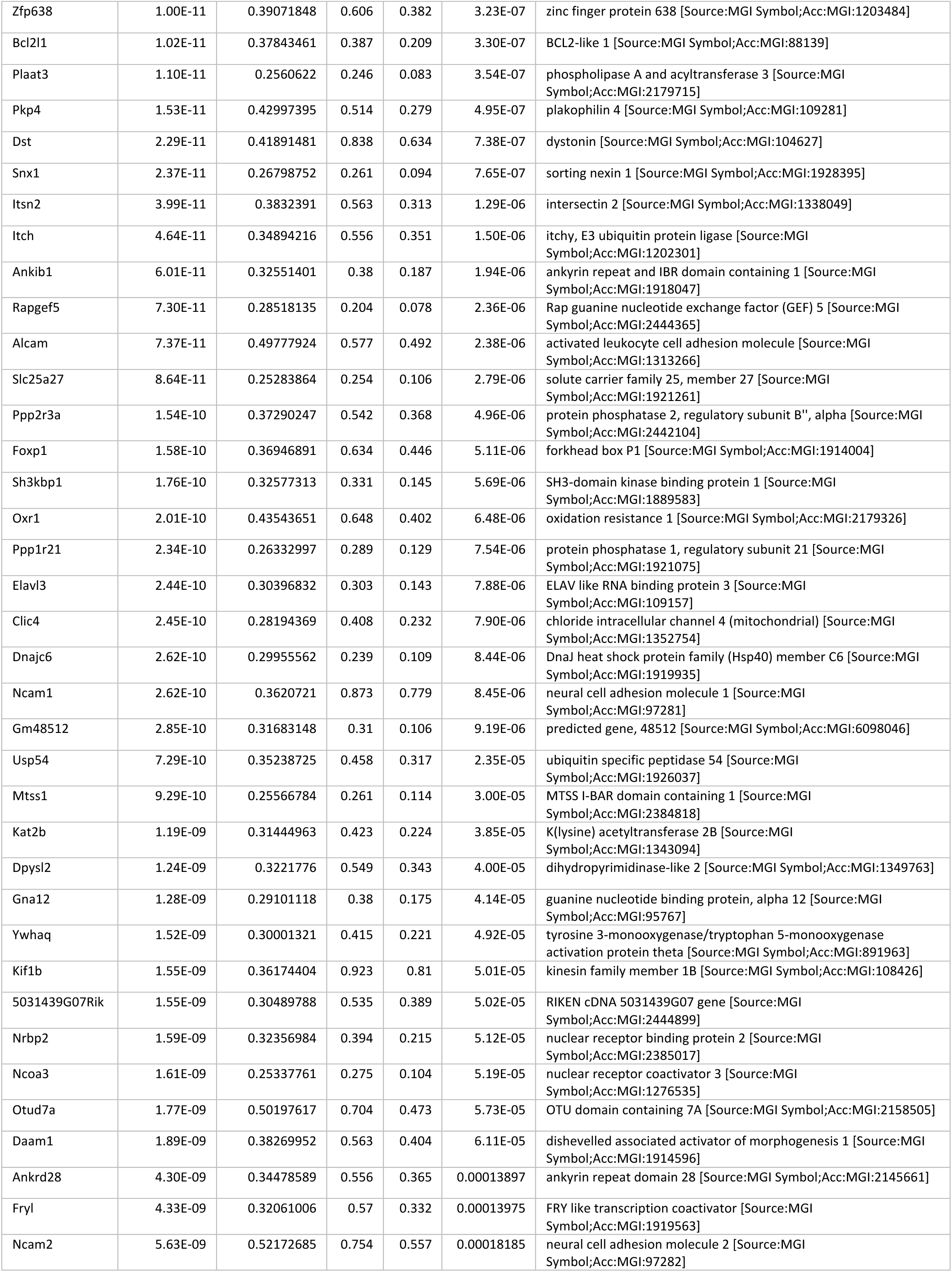
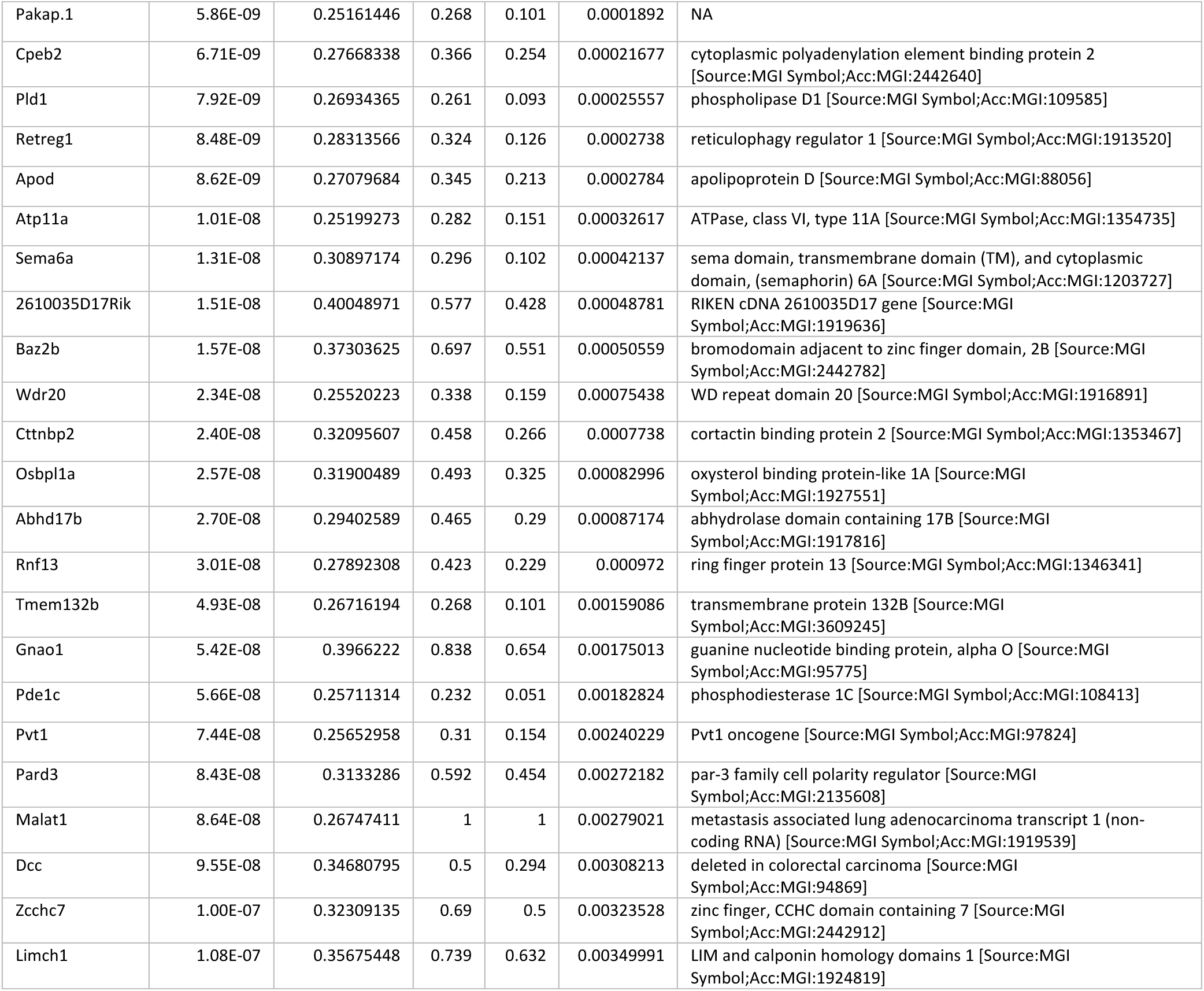
Differentially upregulated genes (p. value < 0.005) in Astrocyte Group 1 in comparison with AG2 and AG3.

Interestingly, Foxp1, a transcription factor with major roles in gene transcription regulation of tissue and cell type-specificity, and involved in subtype identity coordination, migration, and organization of motor neurons (52–54), showed a diverse expression across all groups, but it was significantly upregulated in AG1 of motor cortex. Additional TFs found significantly upregulated included Creb5, Litaf, Nfe2l3, Tulp4, Tcf12, Rere and Pbx3.

Tmem117 transmembrane protein was found expressed uniquely in AG1 from spinal cord, being both group 1 and tissue specific marker.

Achaete-scute family bHLH transcription factor 1 (Ascl1) gene was significantly upregulated in AG1. Magnusson et al. demonstrated that Ascl1+ mature astrocytes take on a progenitor-cell like role and become neurogenic after stroke in adult mice (55). Similar small populations related with progenitor cells have been described in rodent hippocampus, linked with GFAP expression (56), or defined by high expression of Frzb, Ascl1, and Slc1a3 (57), although there is no consensus if misidentification of progenitors in adult brain could be due to de-differentiation of mature neurons (58).

To extend the understanding of these findings in a multicellular scenario and how AG1 might interact with specific sets of neurons, we analyzed cell-cell communication patterns and signaling pathways based on ligand-receptor interactions using the motor cortex expression data from the tree mice, between neuron clusters and the three different AG. Results show SEMA6 signaling and communication pathway exclusively active between AG1 and specific neuron clusters 3, 4 and 8 (Fig. 4c,d,e). SEMA6 signaling network, with known roles in CNS development, neuron migration and axon guidance (59–61), was shown to be active among AG1 with a signaling sender role mediated by expression of Sema6a in AG1 (Fig. 4c,e). All AG1, AG2 and AG3 showed moderated expression of receptors Nrg1 and Nrg3, but Erbb4 receptor was exclusively expressed in AG1. AG3 had exclusively a sender role in the NRG signaling pathway, AG1 on the contrary had a receiver role. Neuregulins (NRGs) are a large family of growth factors that stimulate ERBB receptor tyrosine kinases and are critical for the assembly of the GABAergic circuitry including interneuron migration, neural differentiation, axon and dendrite development, myelination and synapse formation (62–65).

These gene markers and findings have been previously described in scientific literature, in which they have been linked with mature astrocytes promoting a neurogenic environment or acquiring progenitor-cell like properties (55,66–71).

### GFAP-positive astrocyte population

Astrocyte group 2 (AG2) yielded 223 genes that were differentially upregulated (p-value < 0.005, Table 2). Within this list, Gfap was among the top, particularly in the motor cortex, where it was almost exclusively expressed in AG2 cell clusters. Additional markers of astrocyte activation (44) were found upregulated on the AG2 population, including Aldoc, Stat3 and C3.

**Table 2.**
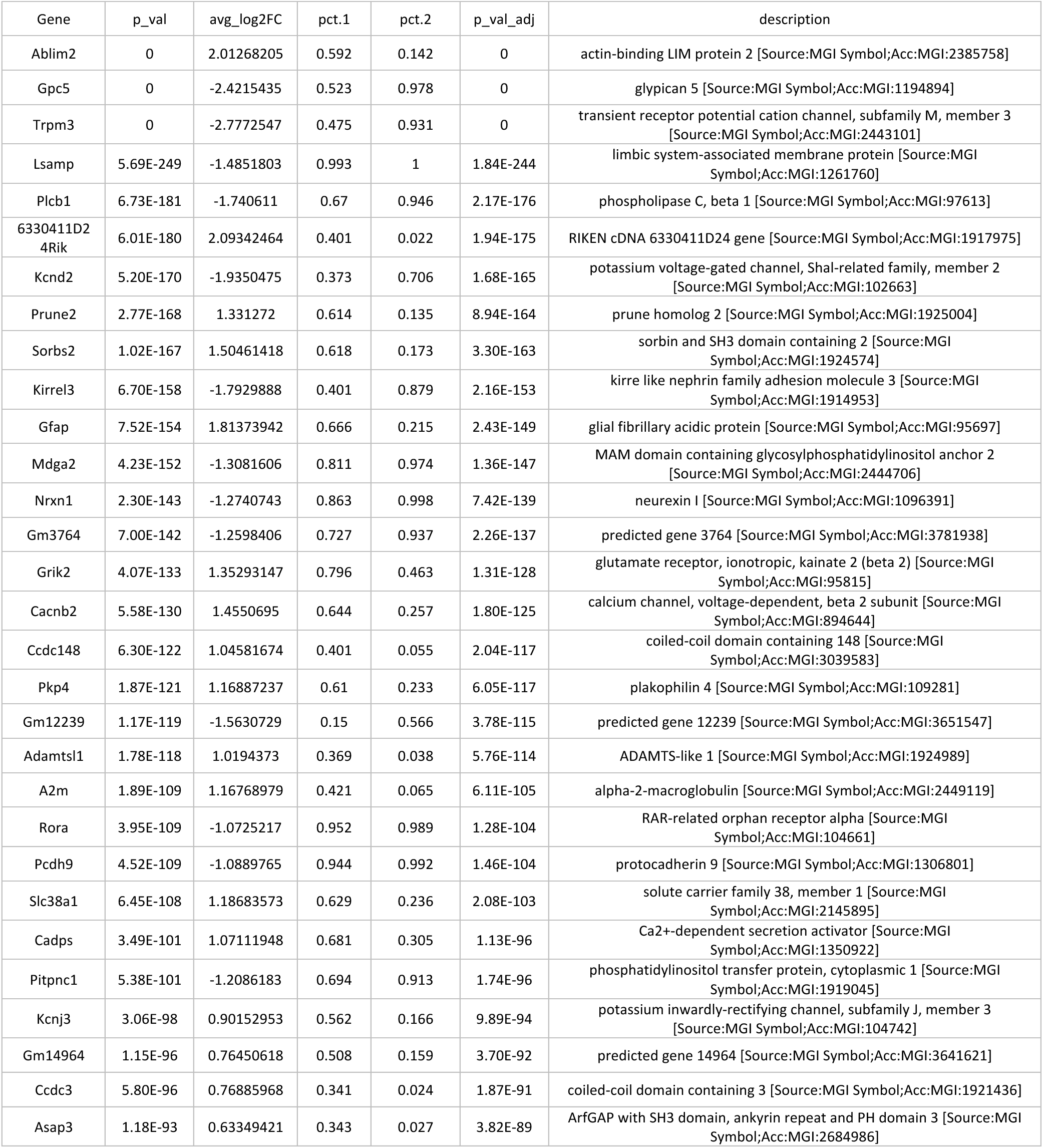
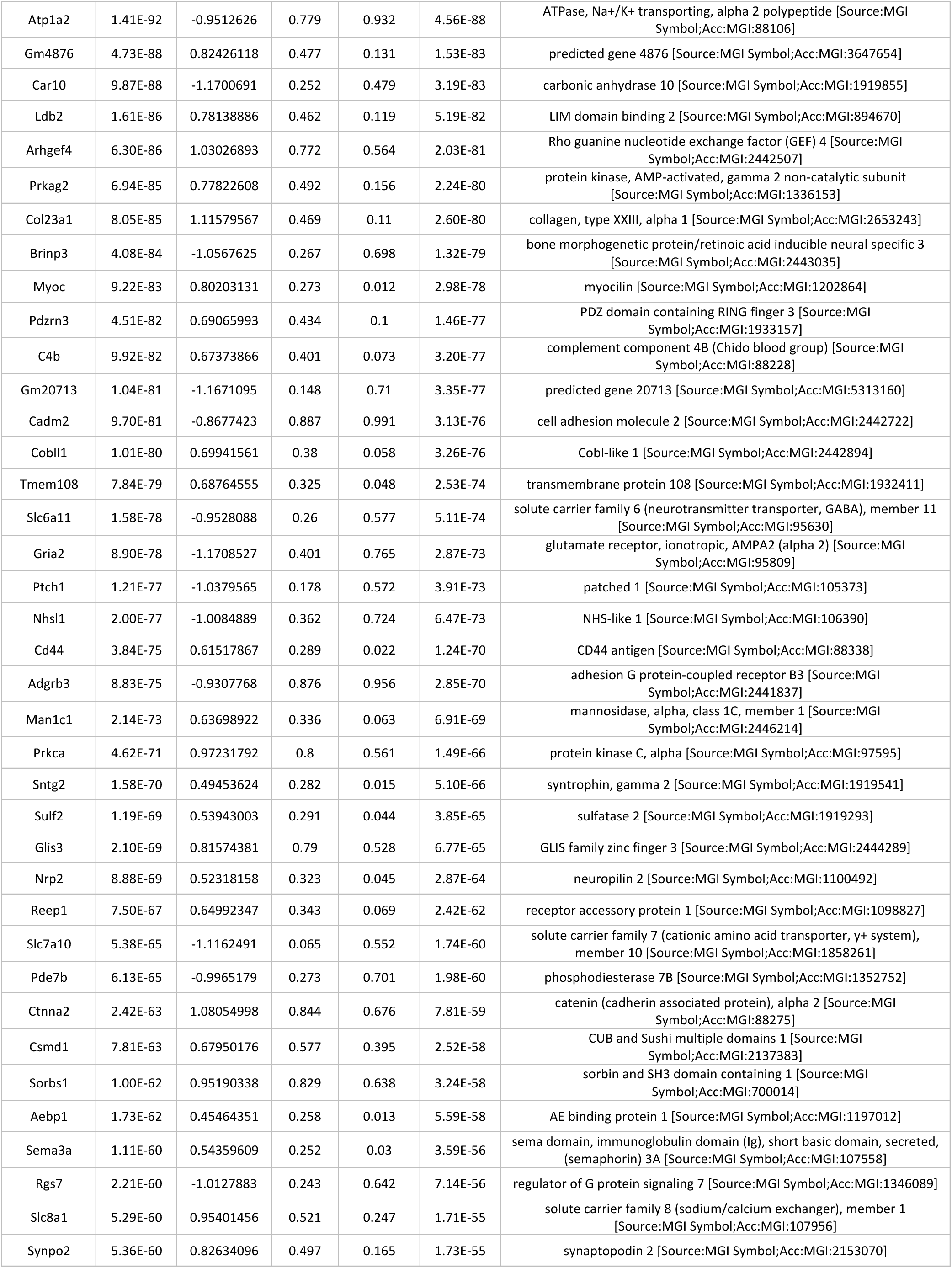
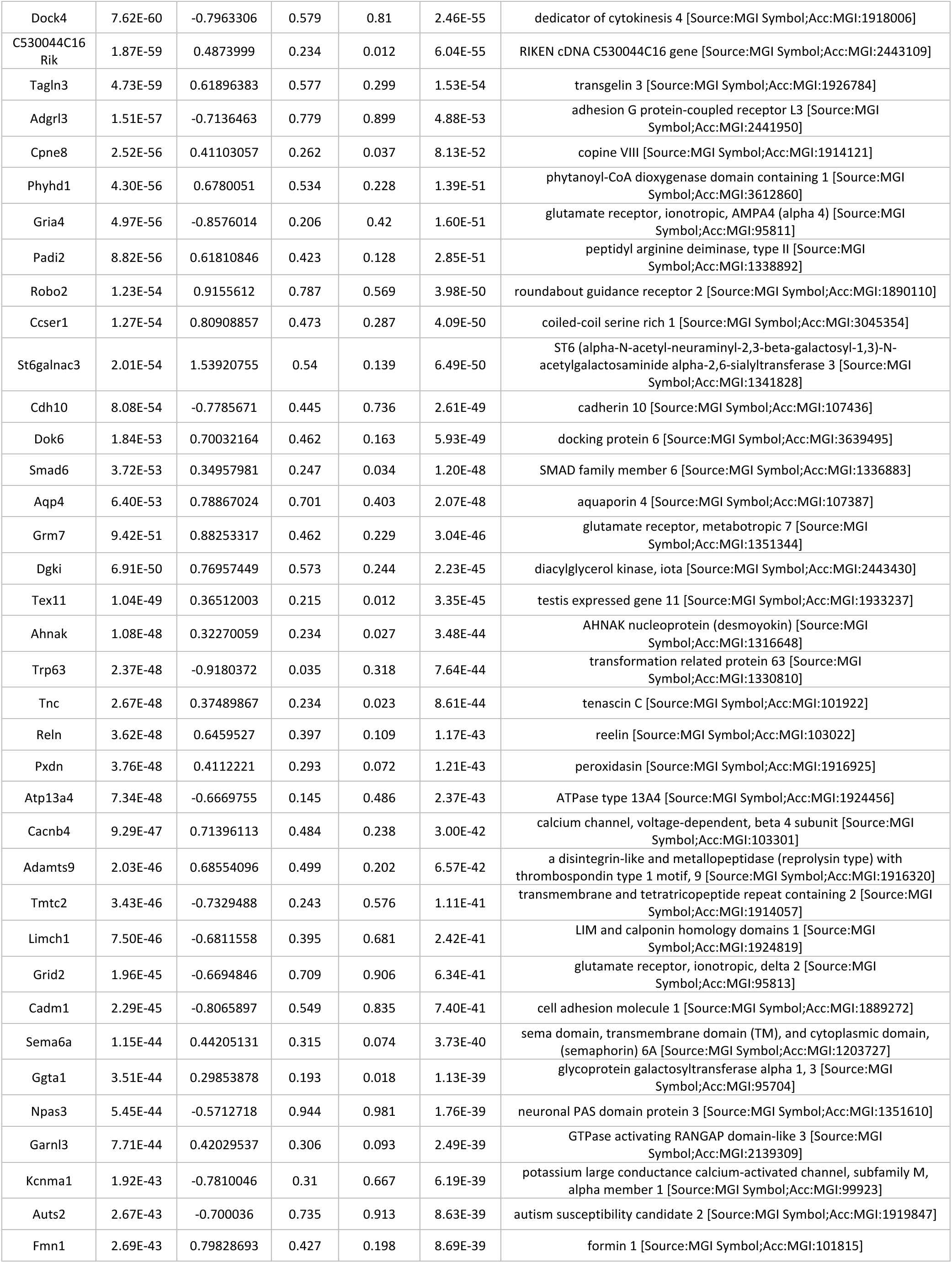
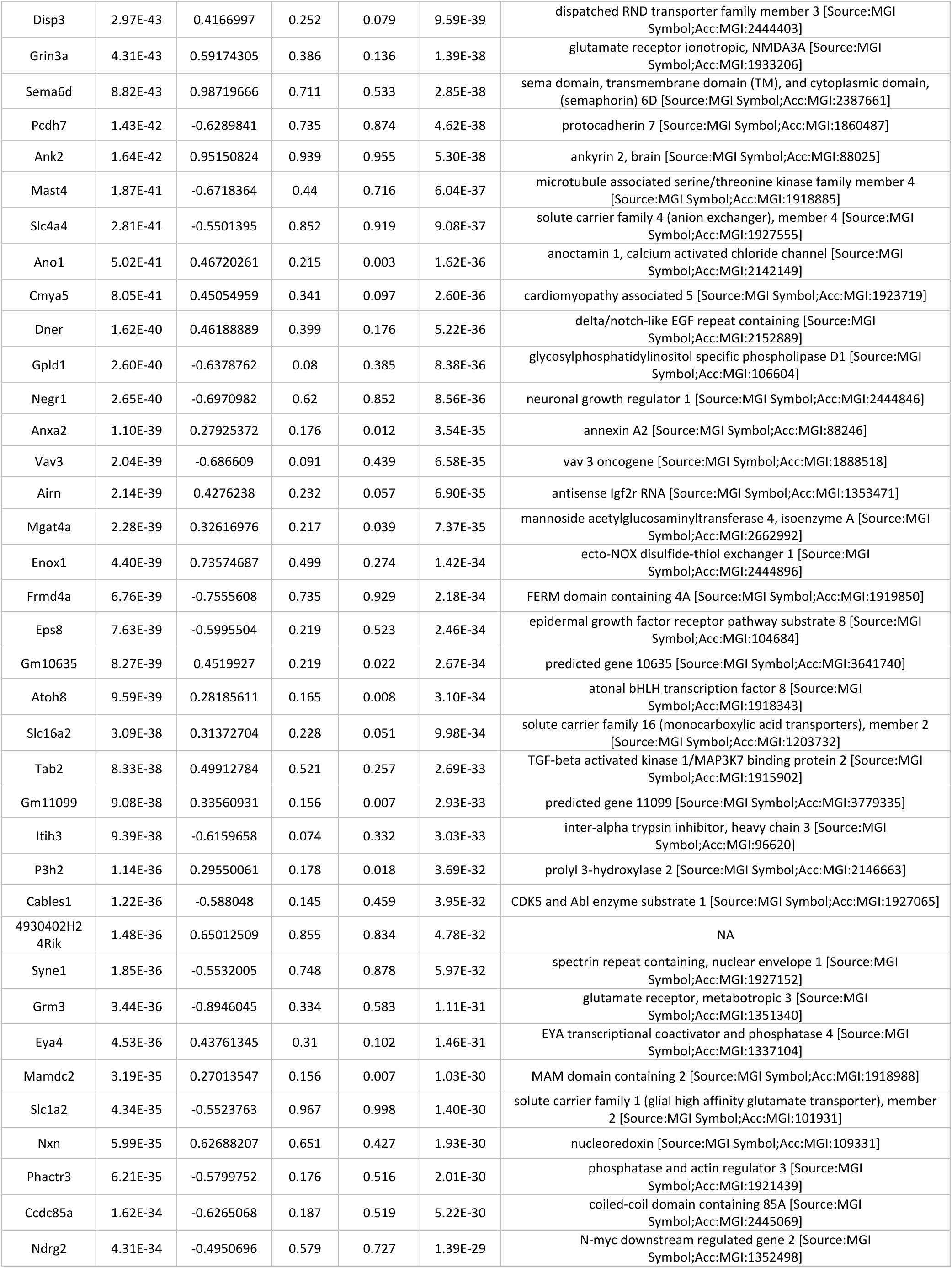
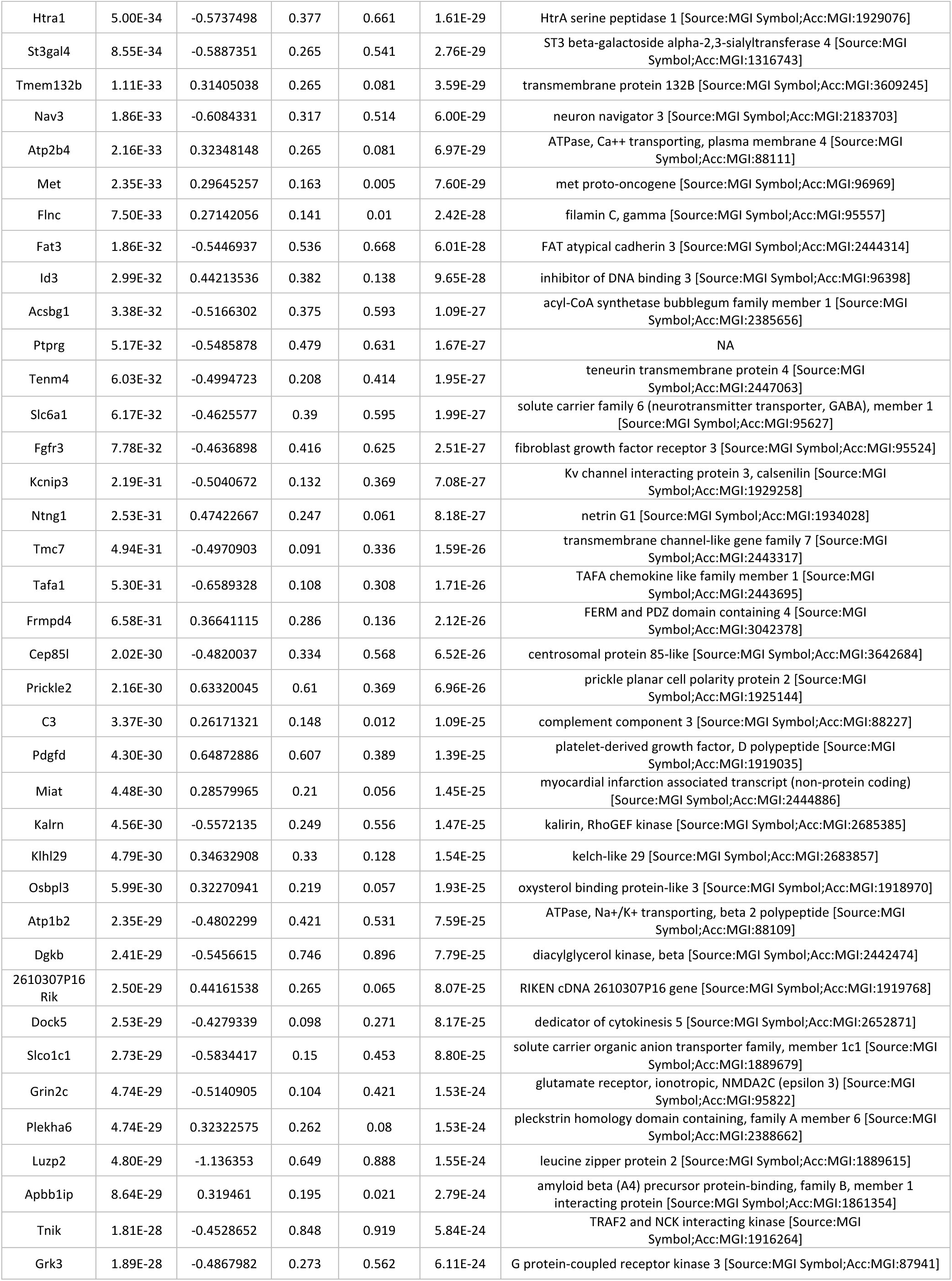
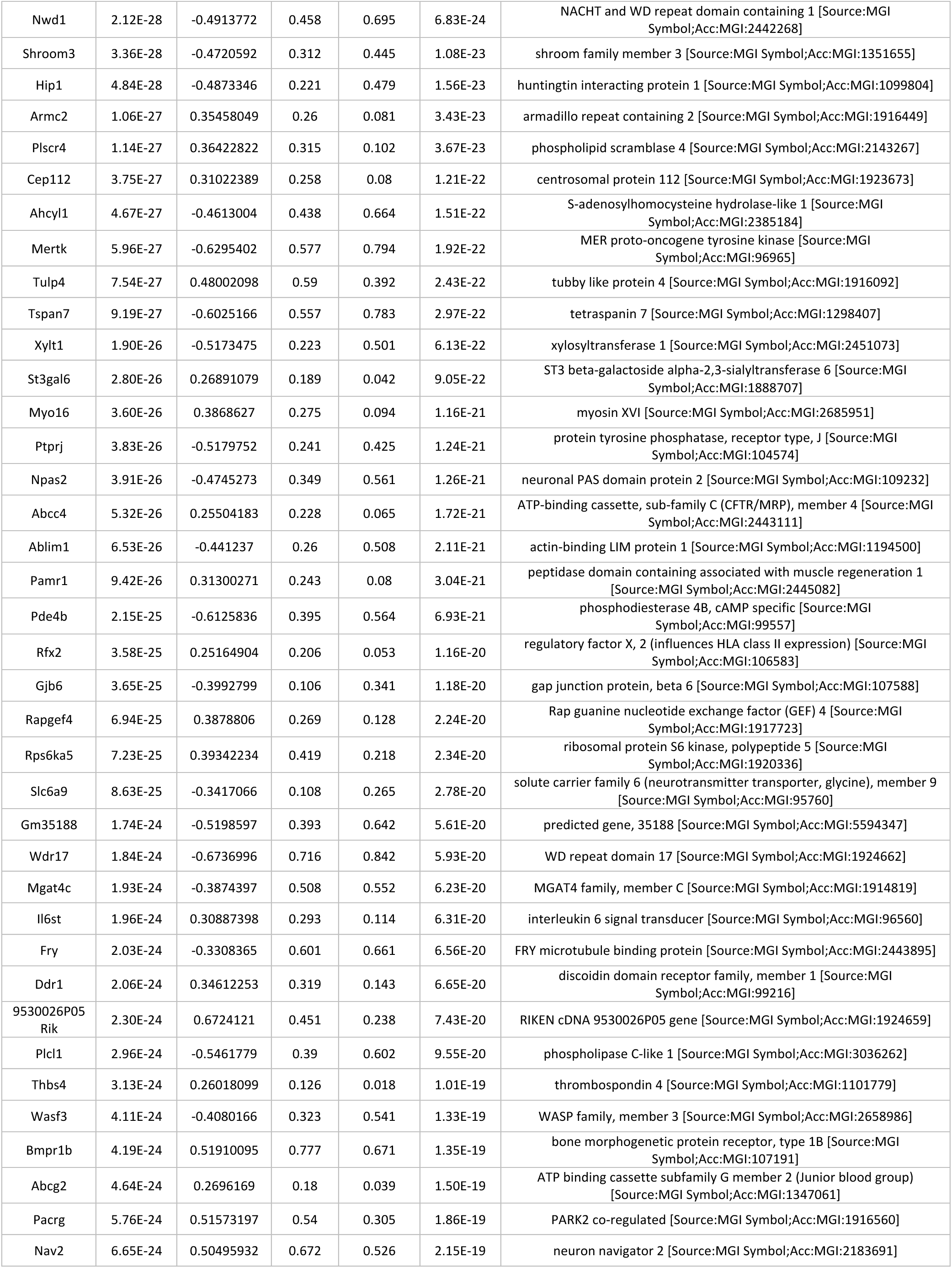
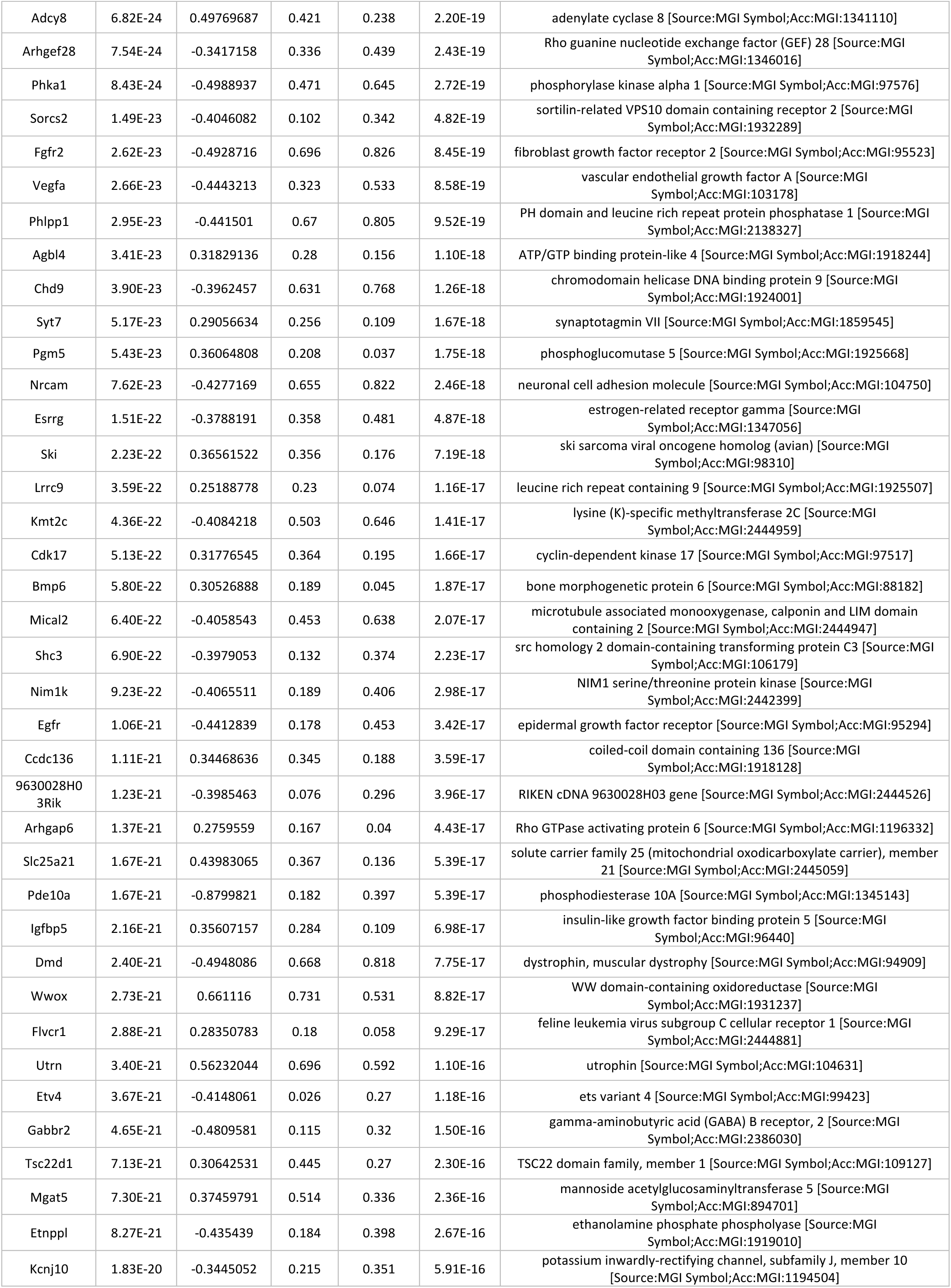
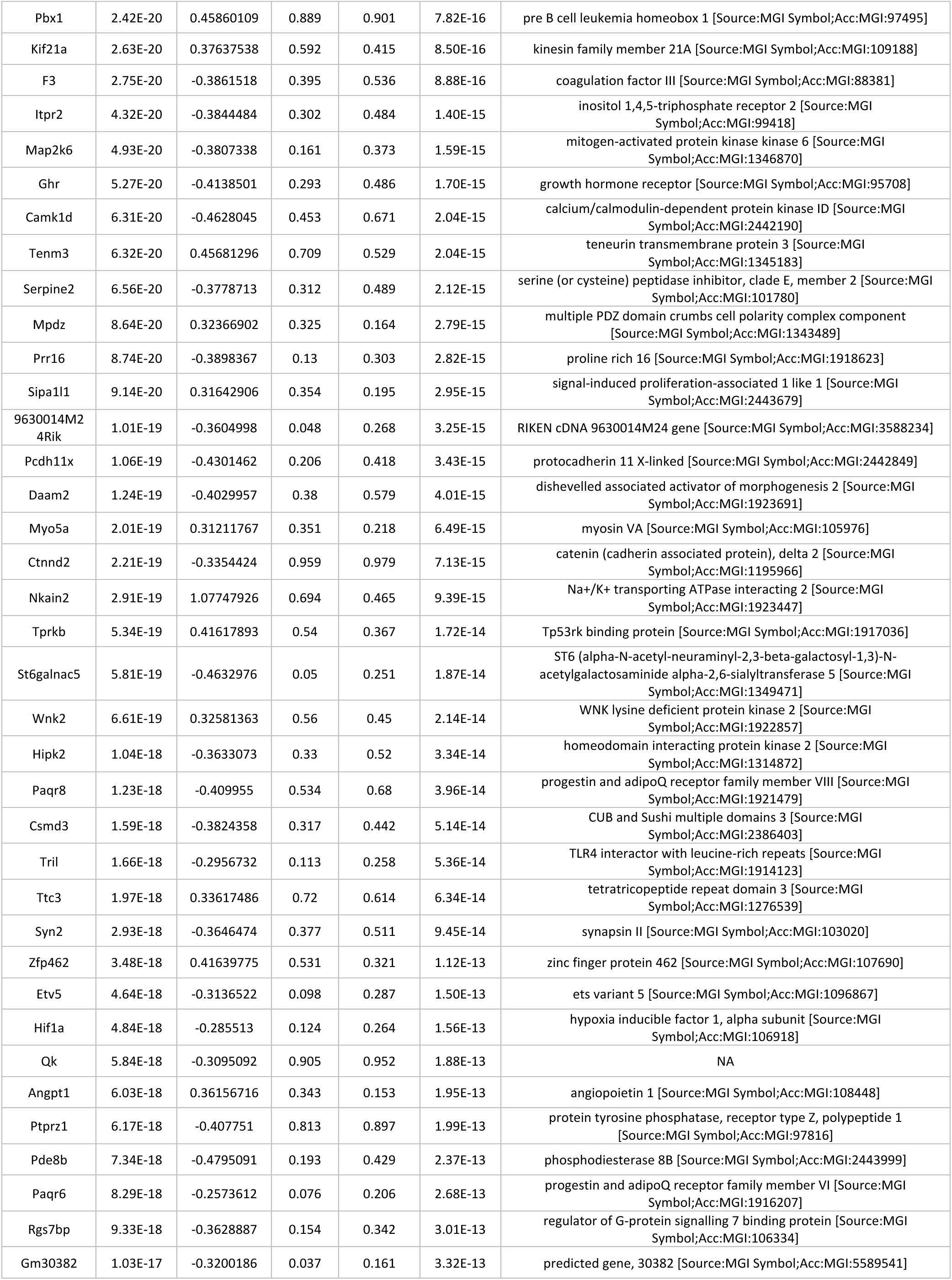
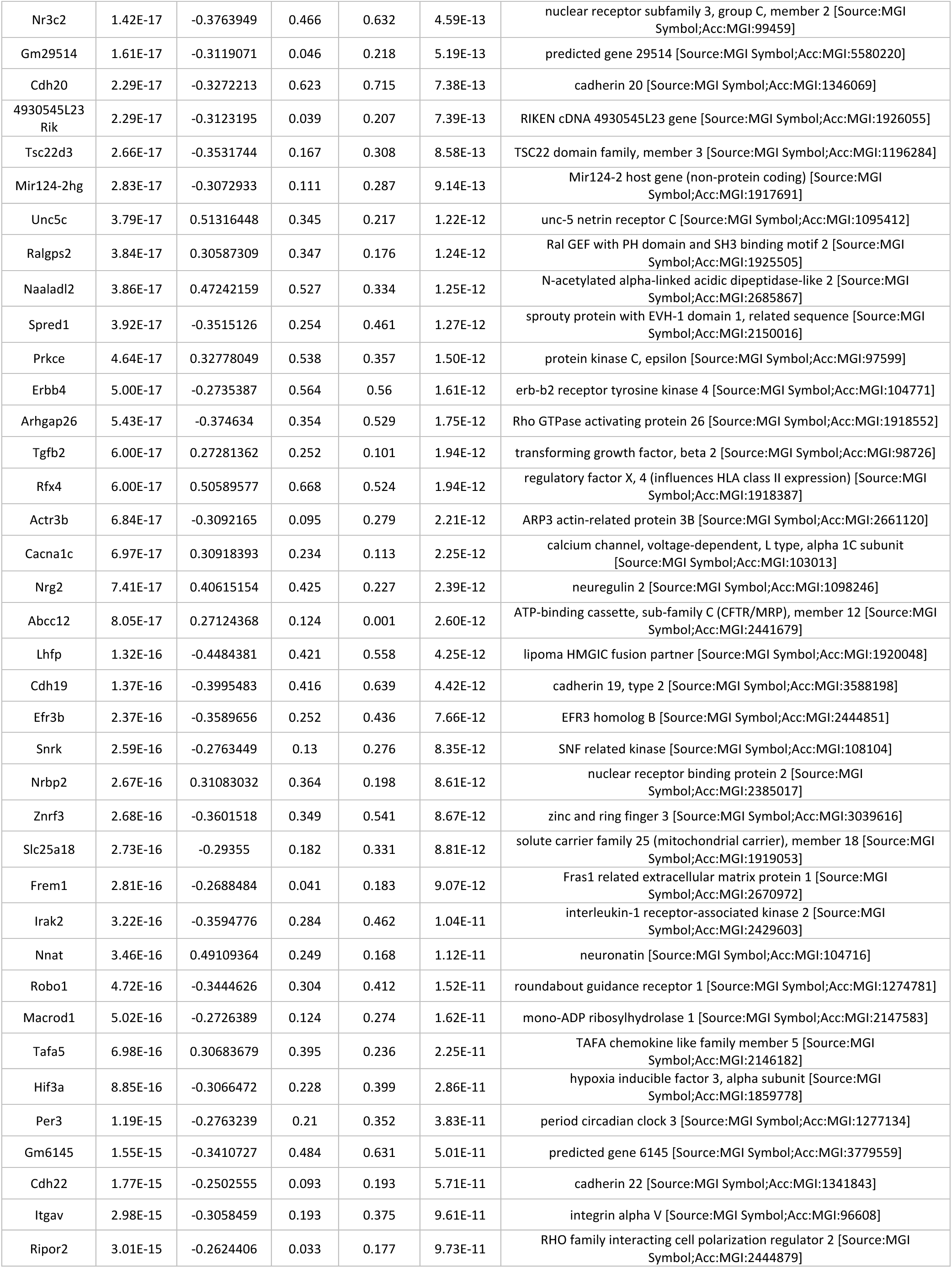
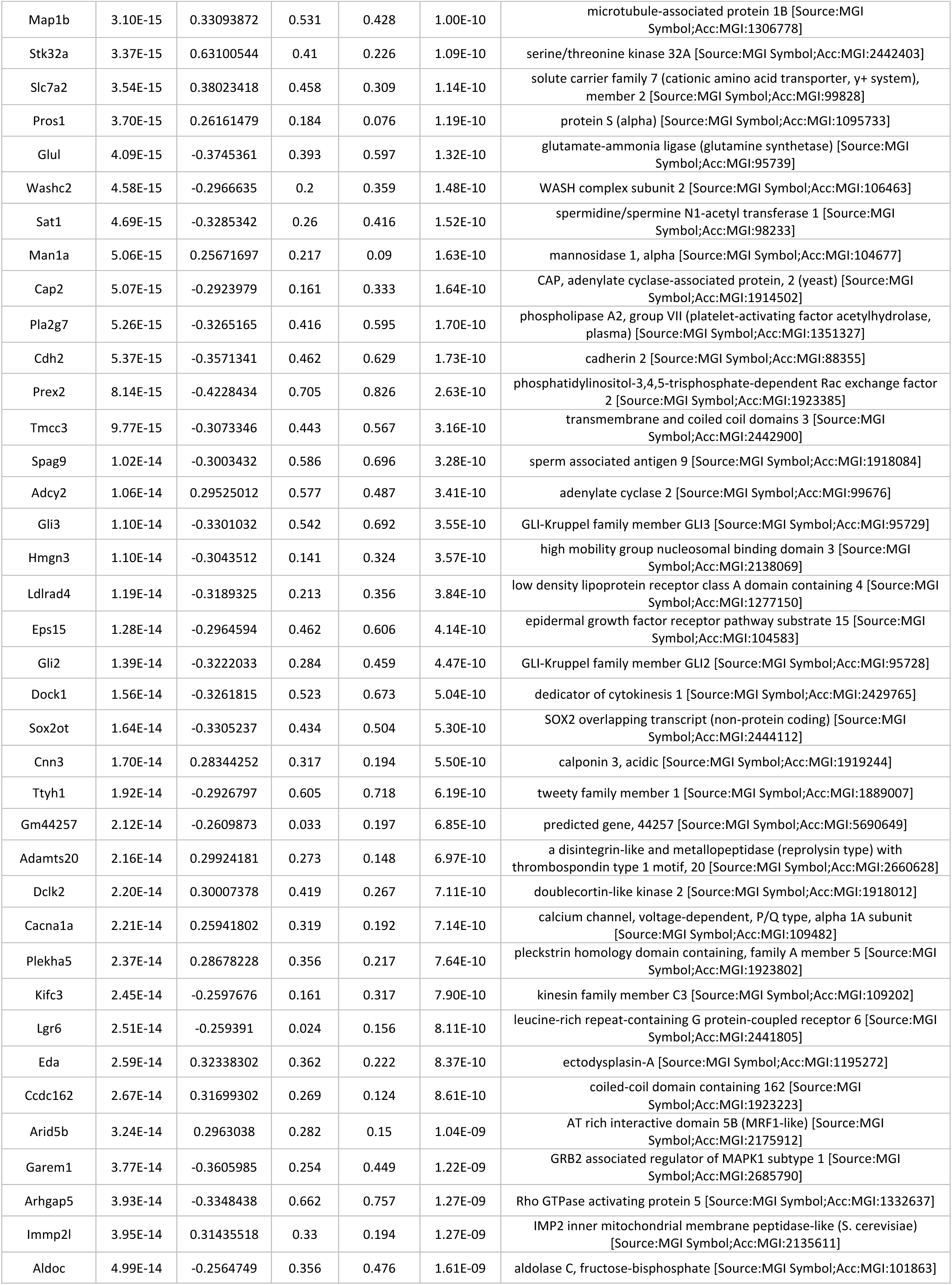
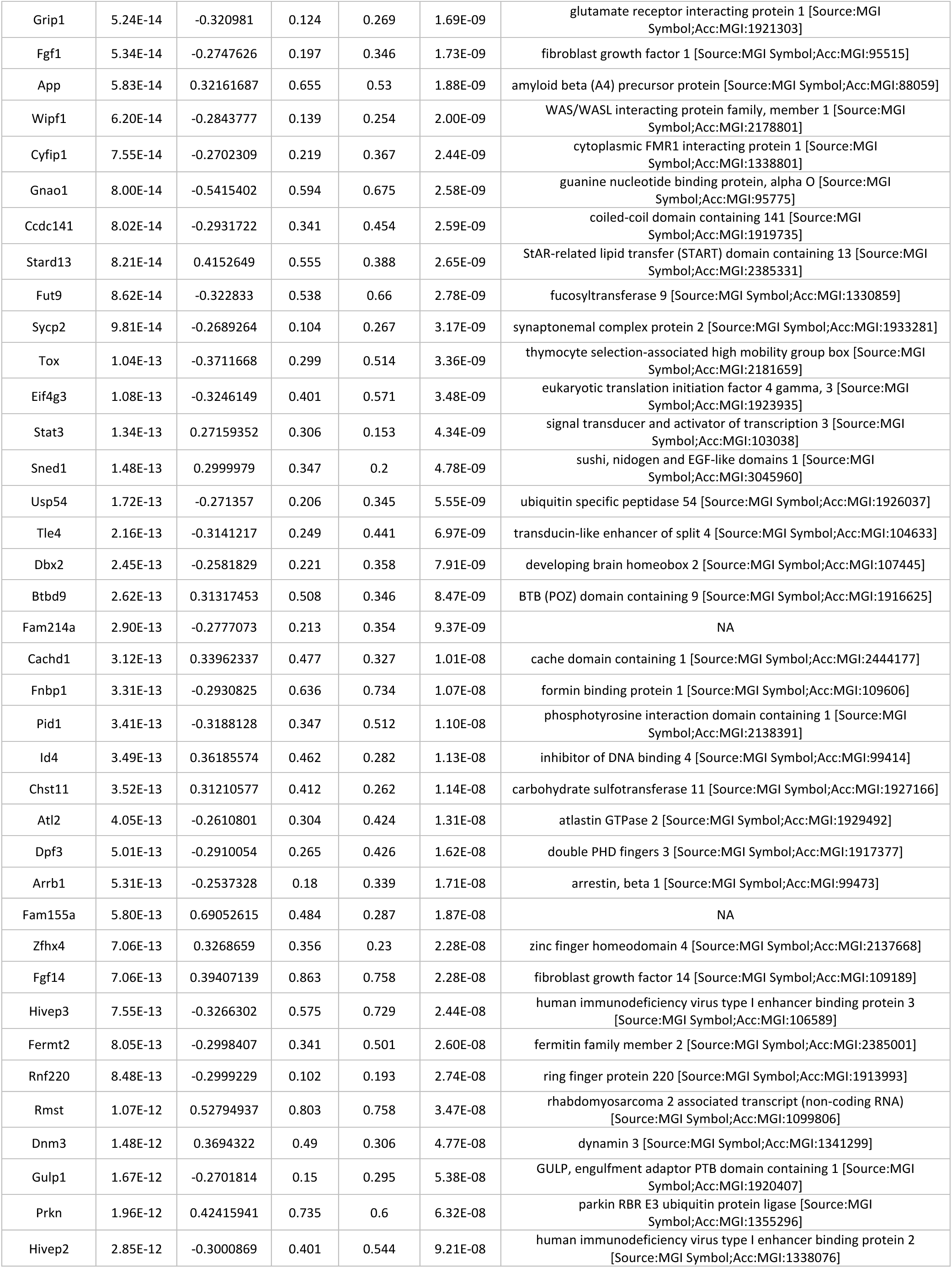
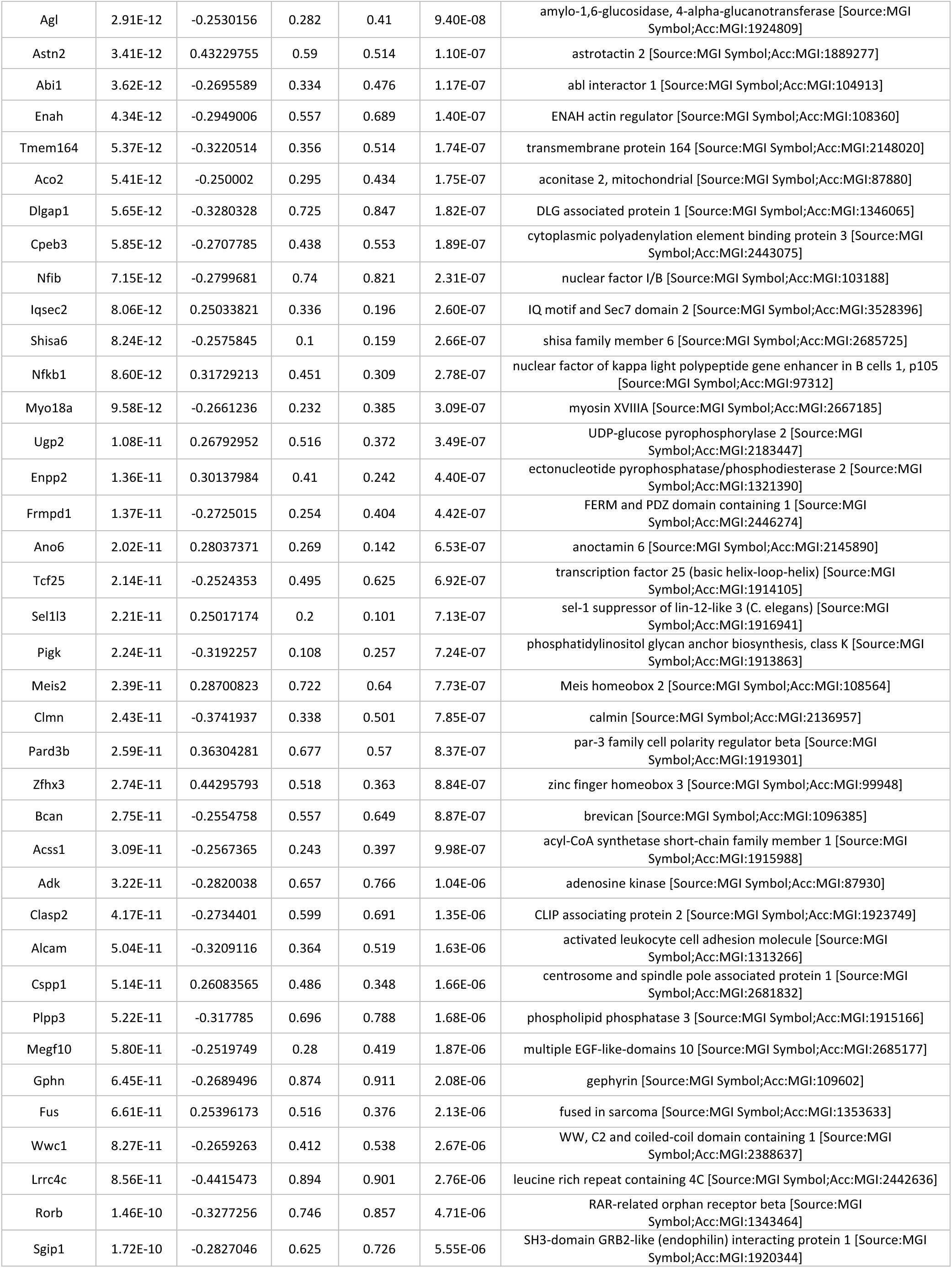
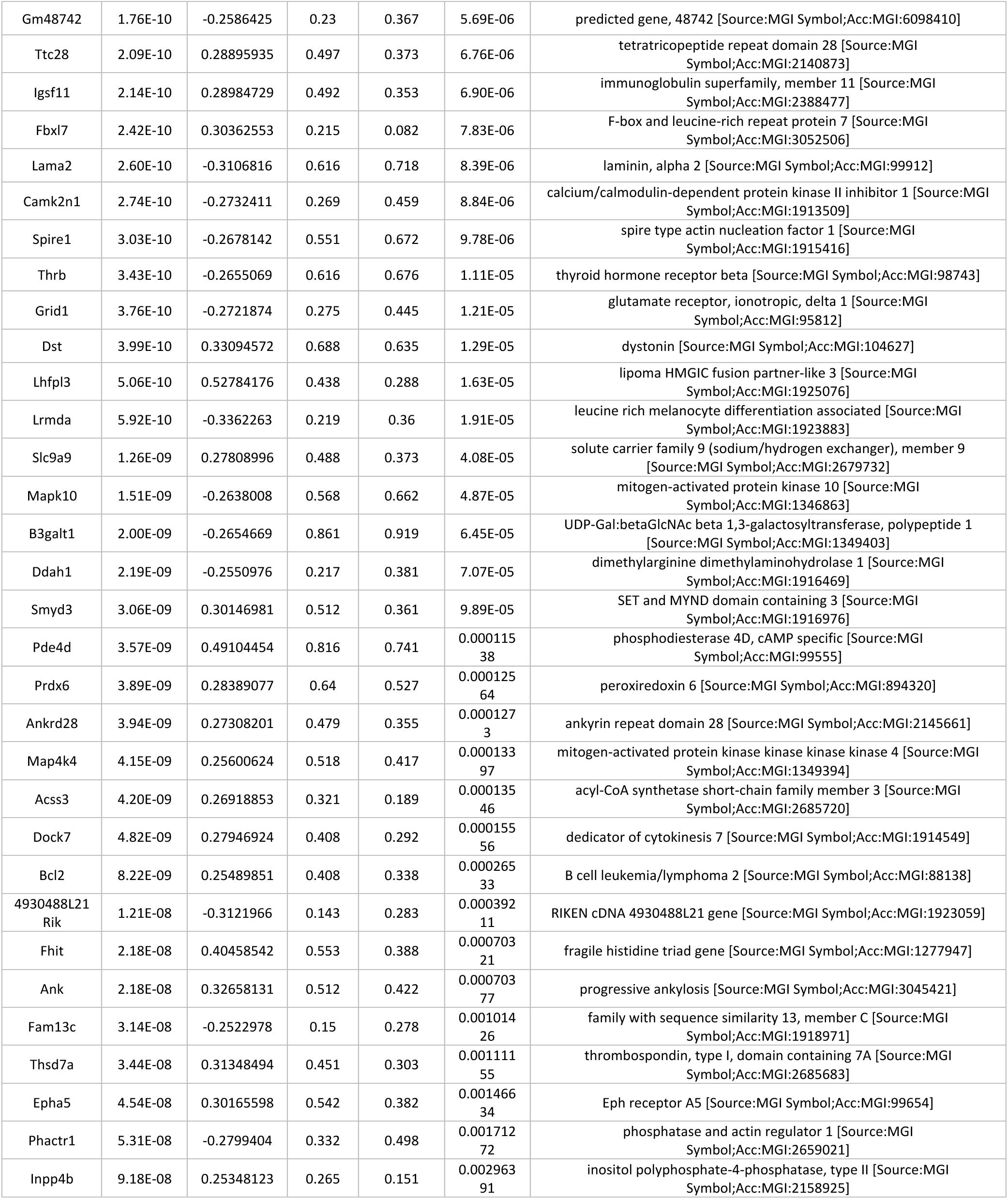
Differentially upregulated genes (p. value < 0.005) in Astrocyte Group 2 in comparison with AG1 and AG3.

Among the enriched biological terms associated with the upregulated genes are neuron and cell differentiation, regulation of cell migration, cell junction organization, and protein binding. Additionally, enriched biological functions unique to AG2 and not observed in AG1 included astrocyte end-foot, T-tubule, actin filament organization, and regulation of cell migration.

Moreover, we identified 16 transcription factors (only 1 shared with AG1 population) found upregulated, including Glis3, Stat3, Smad6, Atoh8, Arid5b, Nfkb1 and Id3.

Complement component 3 (C3) was exclusively expressed in AG2 of spinal cord. GFAP/C3-positive astrocytes subpopulations have been reported in numerous brain samples from neurodegenerative disease patients such as Alzheimer’s disease, multiple sclerosis, Huntington’s disease, Parkinson’s disease, and amyotrophic lateral sclerosis (ALS) (44,72,73).

The expression of Gfap and other accompanying markers links AG2 with the Gfap positive astrocyte population reported in previous studies and described as an activated-state glial population in normal physiological conditions, and reactive astrocytes in disease (41,44,74–76).

### Identification of a large astrocytic population related with mature astrocyte functions

Astrocyte Group 3, by comparison, was the largest and most heterogeneous. Each AG3 was comprised on average by 3 clusters on each sample indicating intra-specimen heterogeneity, with a high similarity score and bound tightly together in the similarity graph (Fig. 3b). We found 158 genes significantly overexpressed (p-value < 0.005) in contrast to AG1 and AG2 (Table 3). Biological term enrichment and ontology analysis of upregulated genes uncover functions such as post synapse, neuron projection, synaptic membrane, glutamatergic synapse, neuron to neuron synapse, postsynaptic specialization, and postsynaptic density.

**Table 3.**
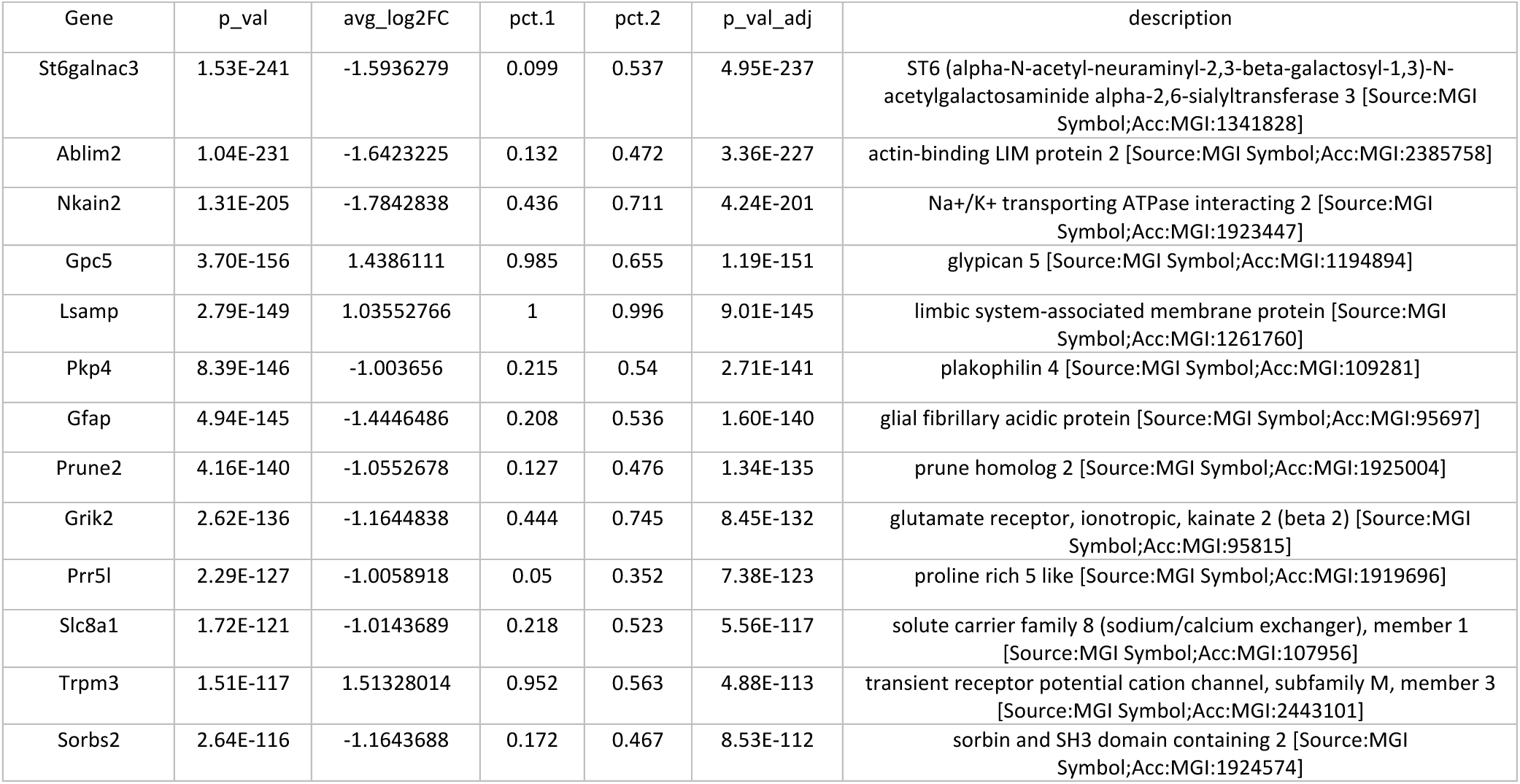
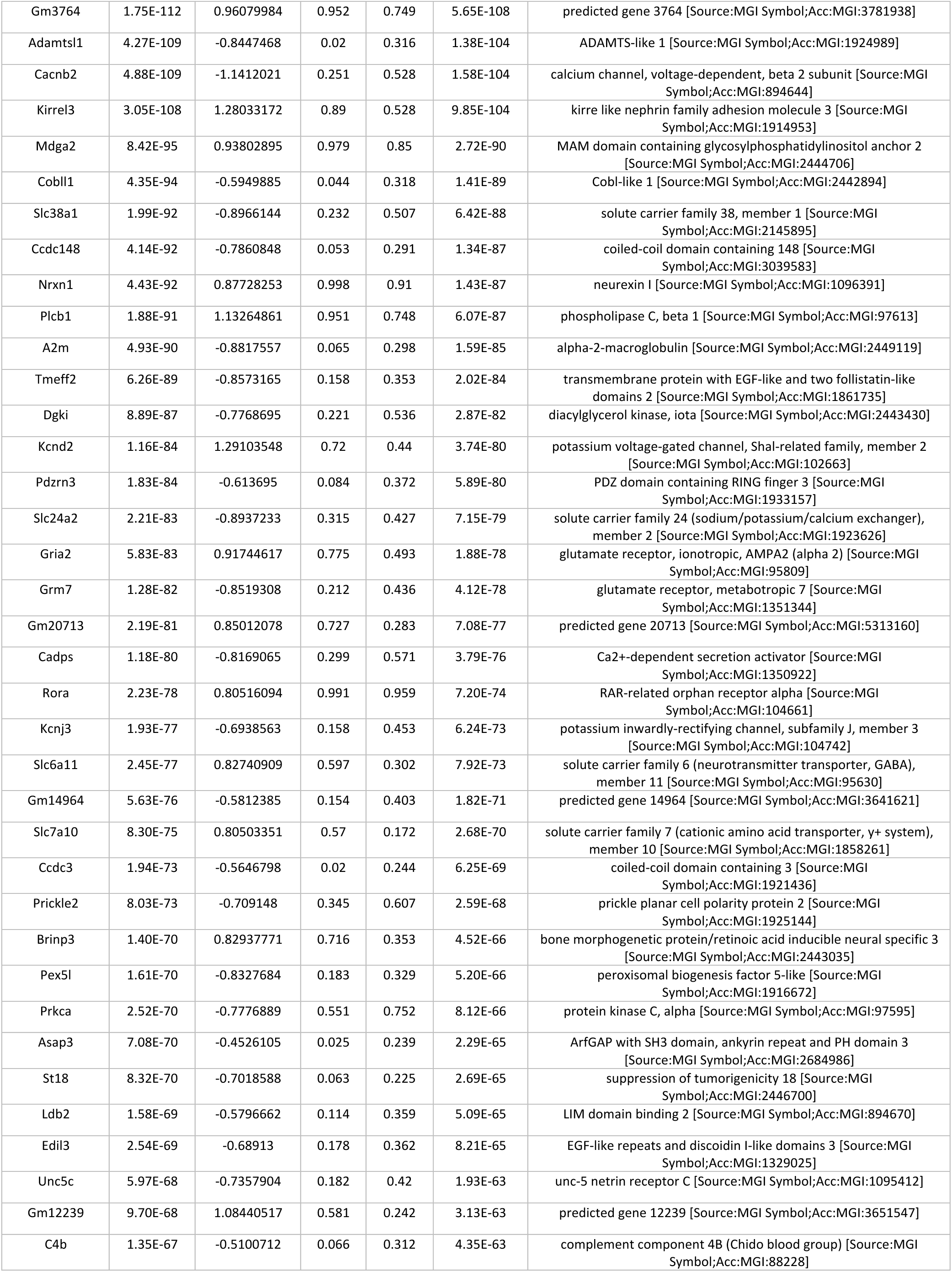
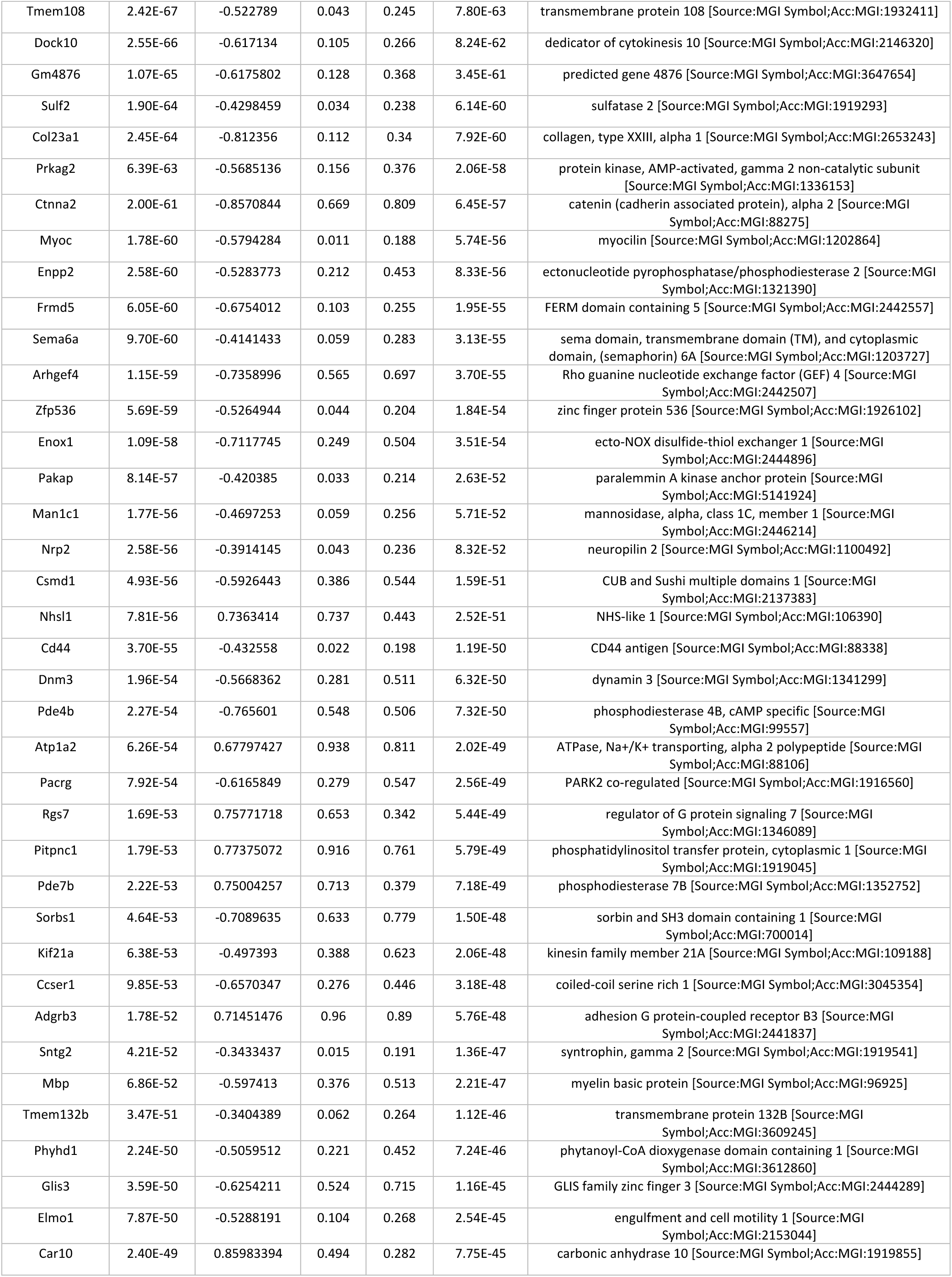
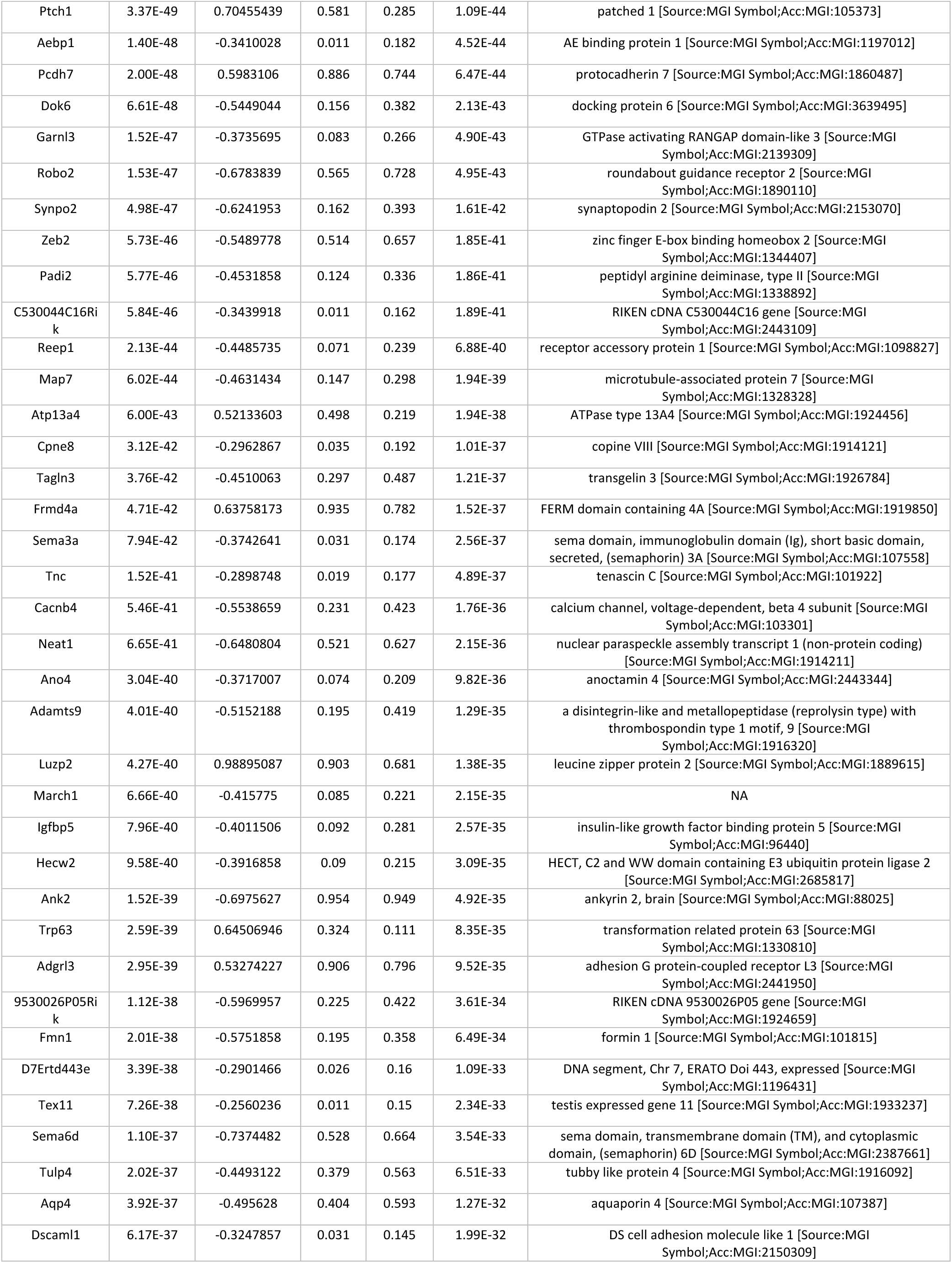
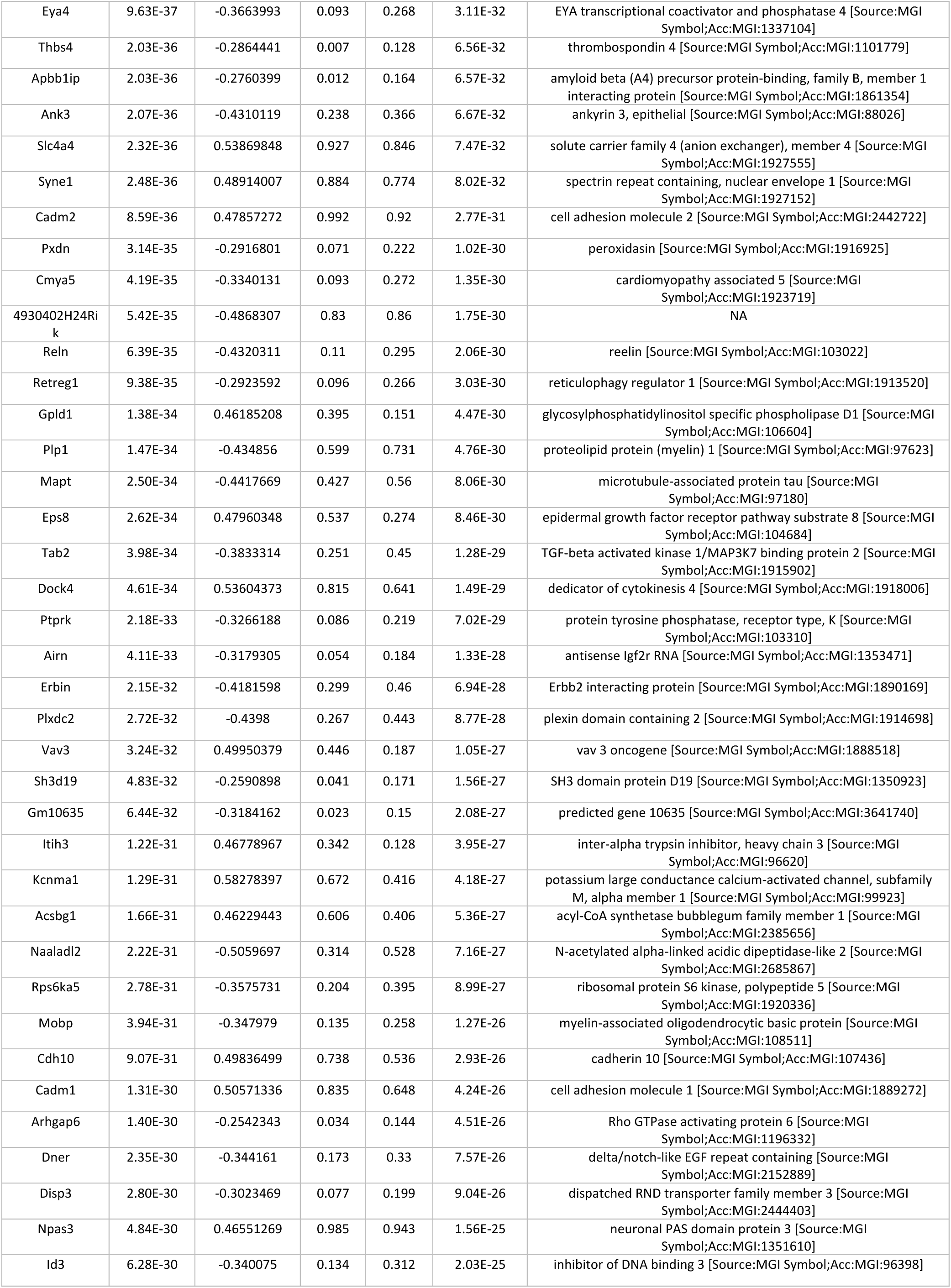
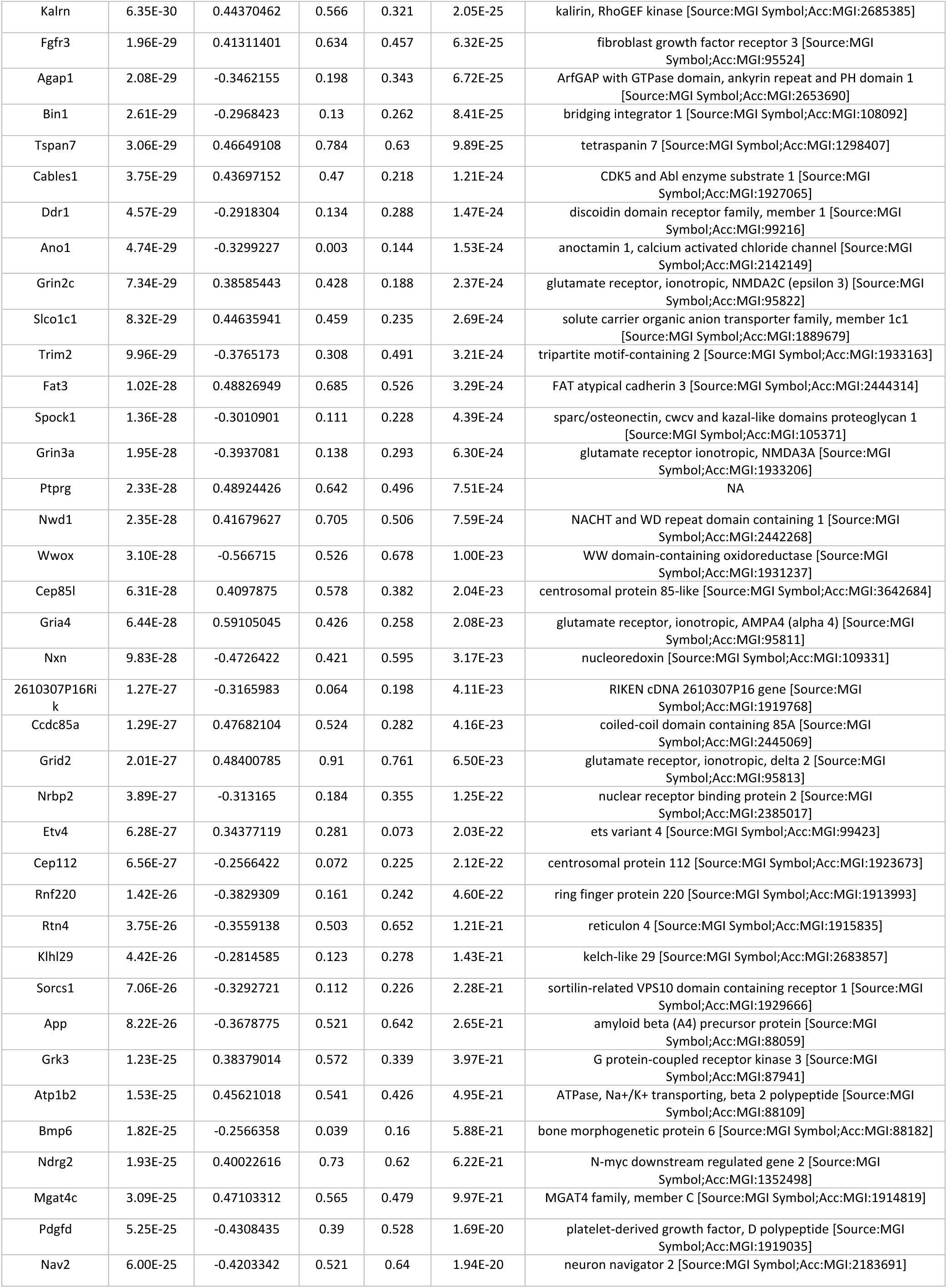
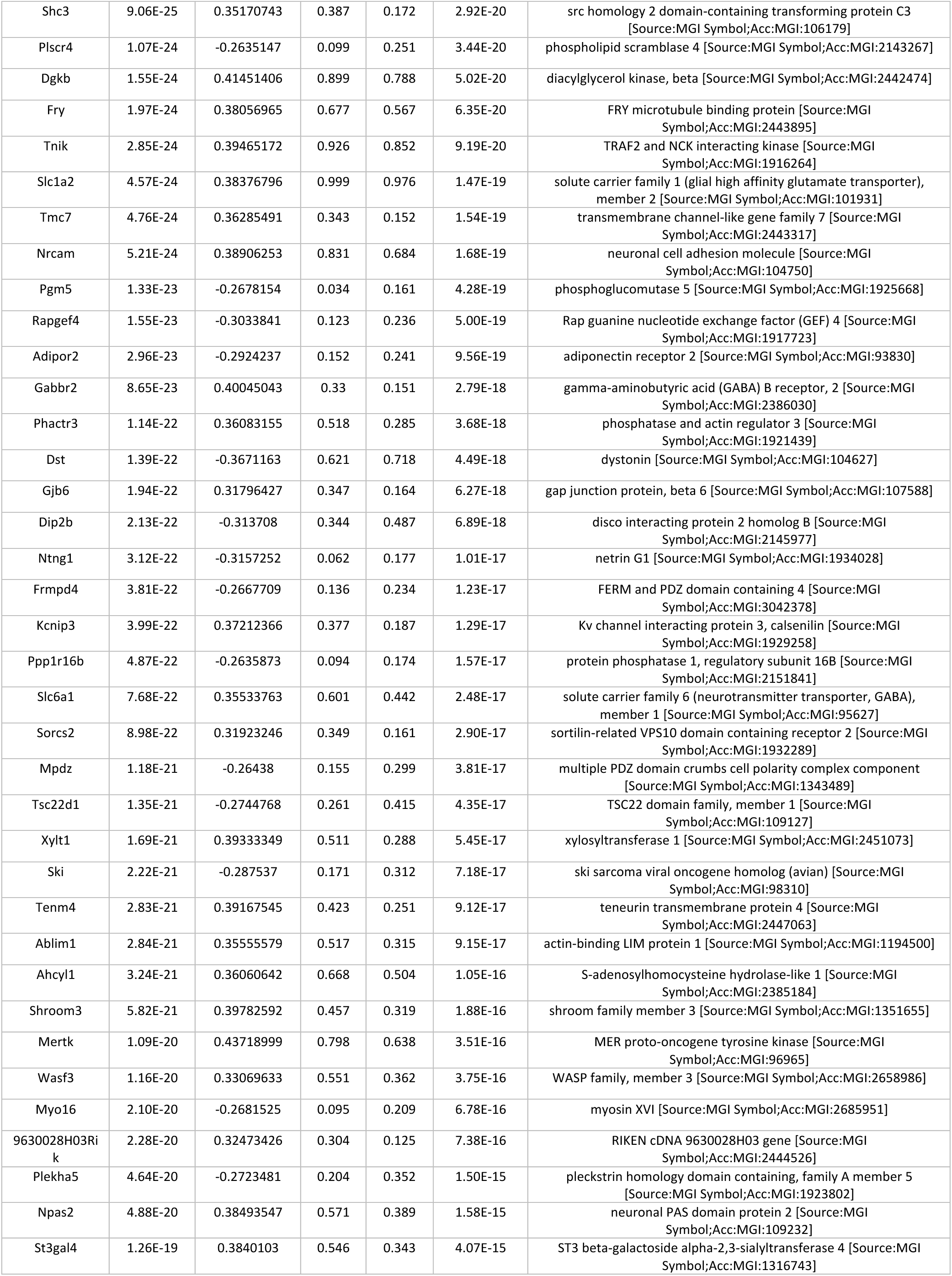
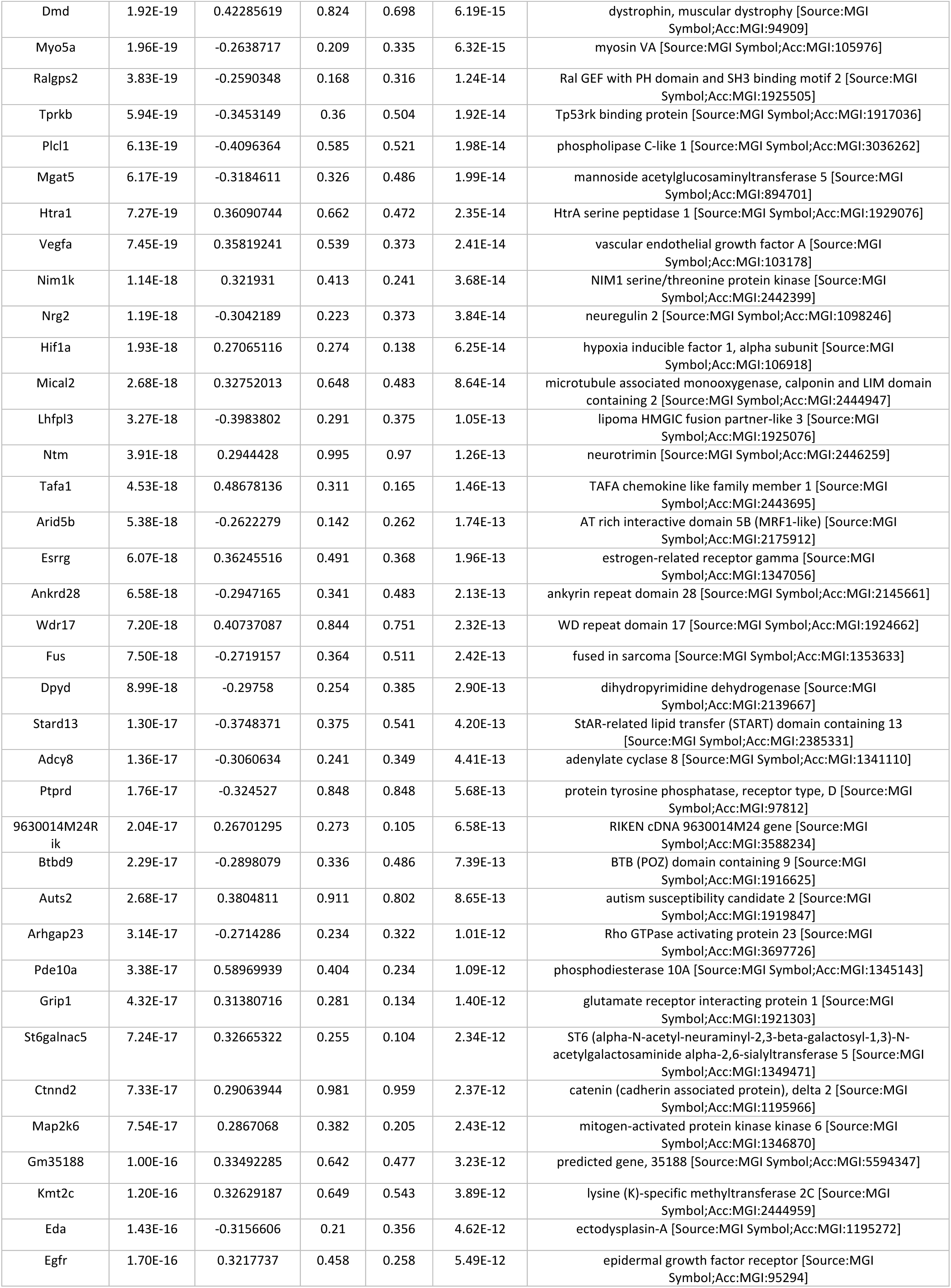
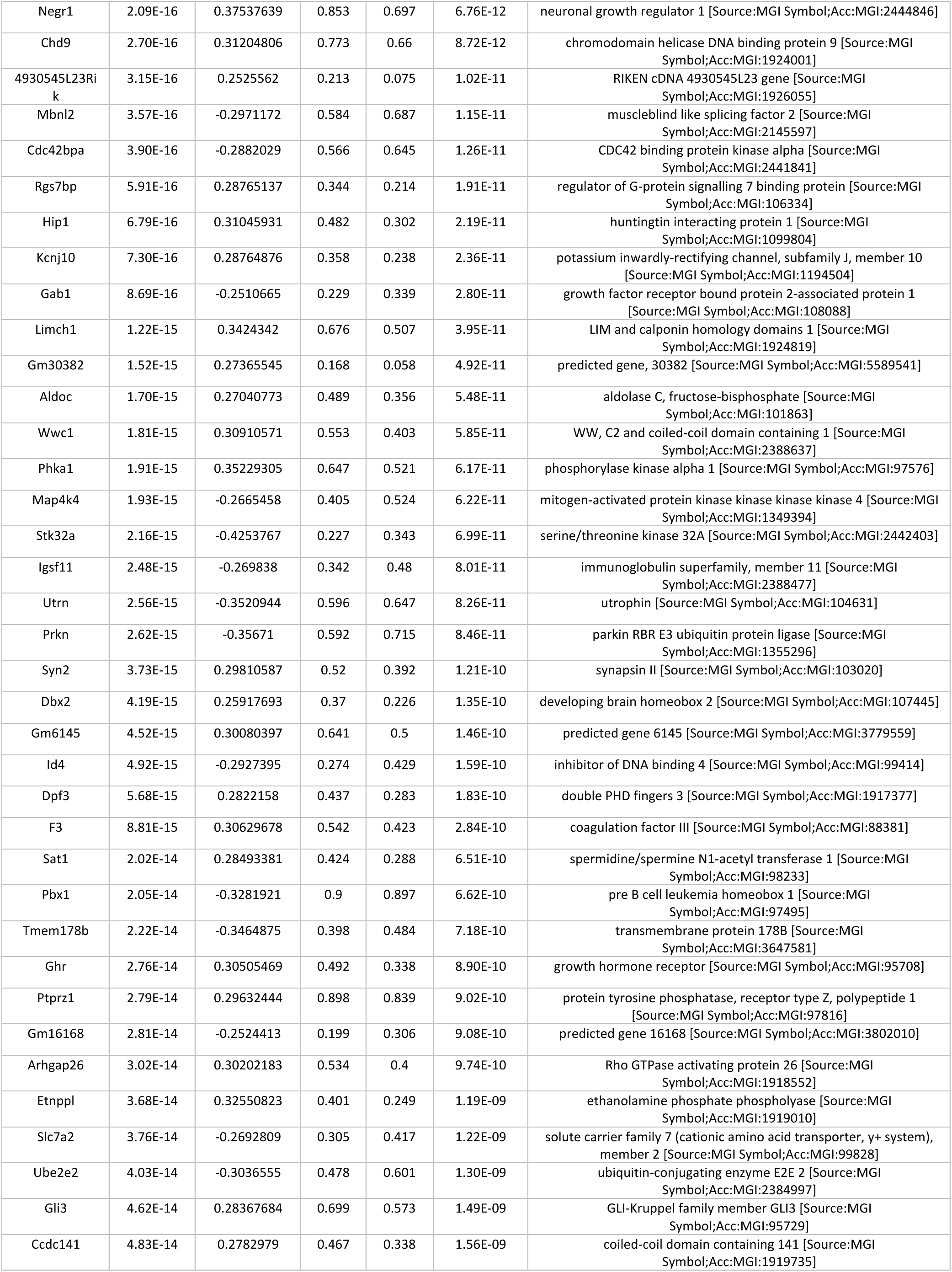
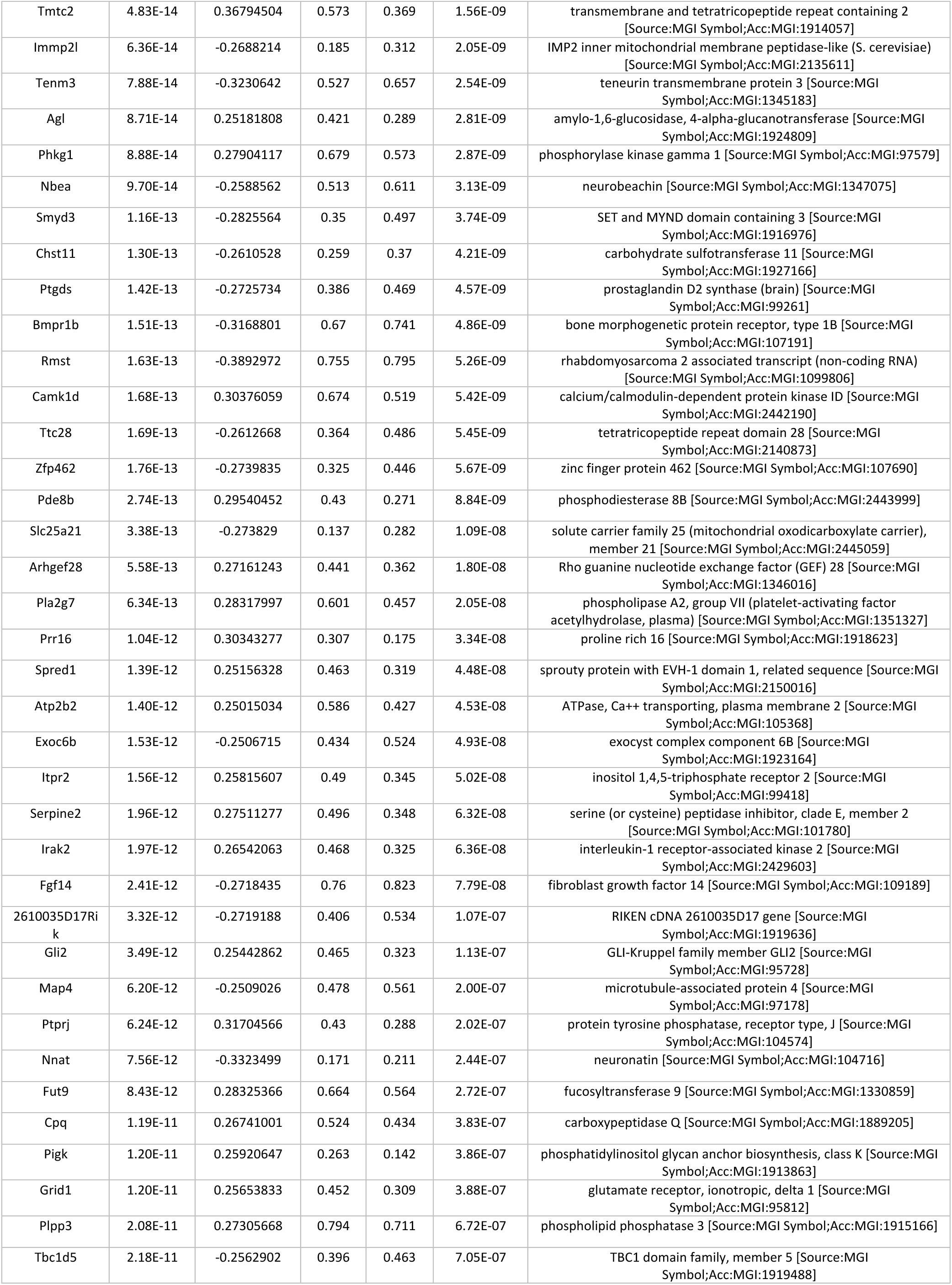
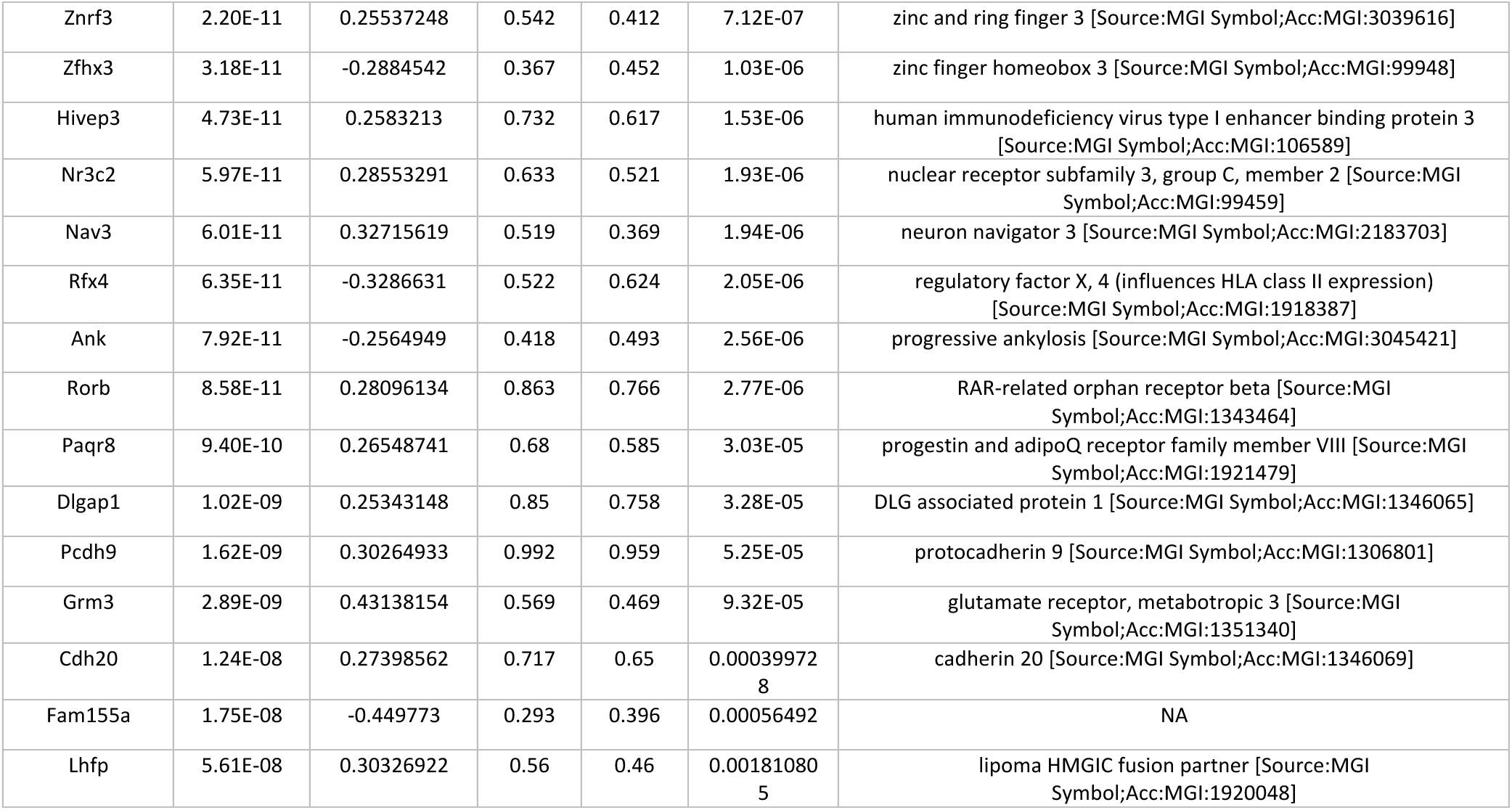
DEG between AG3 and the other astrocyte groups AG1 and AG2.

To interrogate the inner variability of AG3, we performed an individual similarity test using CFS metric by sub-setting the clusters from AG3 cells and performing a community analysis on the resulting graph. Results revealed 3 distinct sub-groups (Fig. 5a), that once annotated, showed similar sources of variability on the merged data of AG3 cells as a second test (Fig. 5b). We decided to include cluster 2 from sample sc.699 on AG3 group 3 as the number of inward edges was higher from subgroup 3 than from the assigned community group 2. We hypothesized these sub-populations corresponded to transient astrocytic states. To test this, we computed pseudotime scores using Monocle2 (77) for these 3 distinct groups using the merged data (Fig. 5b). Top genes found differentially expressed as a function of pseudotime were linked by similar terms related with cell junction, cell-cell and synapse signaling, glutamatergic synapse, and transmembrane transport (Figure 5d). Cell-cell communication analysis using the motor cortex expression data from the tree mice found CNTN and NCAM signaling pathways exclusively active between AG3 and neuronal subclusters (Fig. 5c). Contactin (CNTN) family of cell recognition molecules are involved in the maintenance of the nervous system (78). Neural Cell Adhesion Molecule (NCAM) mediates neural cell–cell interactions and neurite outgrowth (79) and facilitates the interaction between astrocytes and synapses, modulating neuronal communication (80).

**Figure 5.**
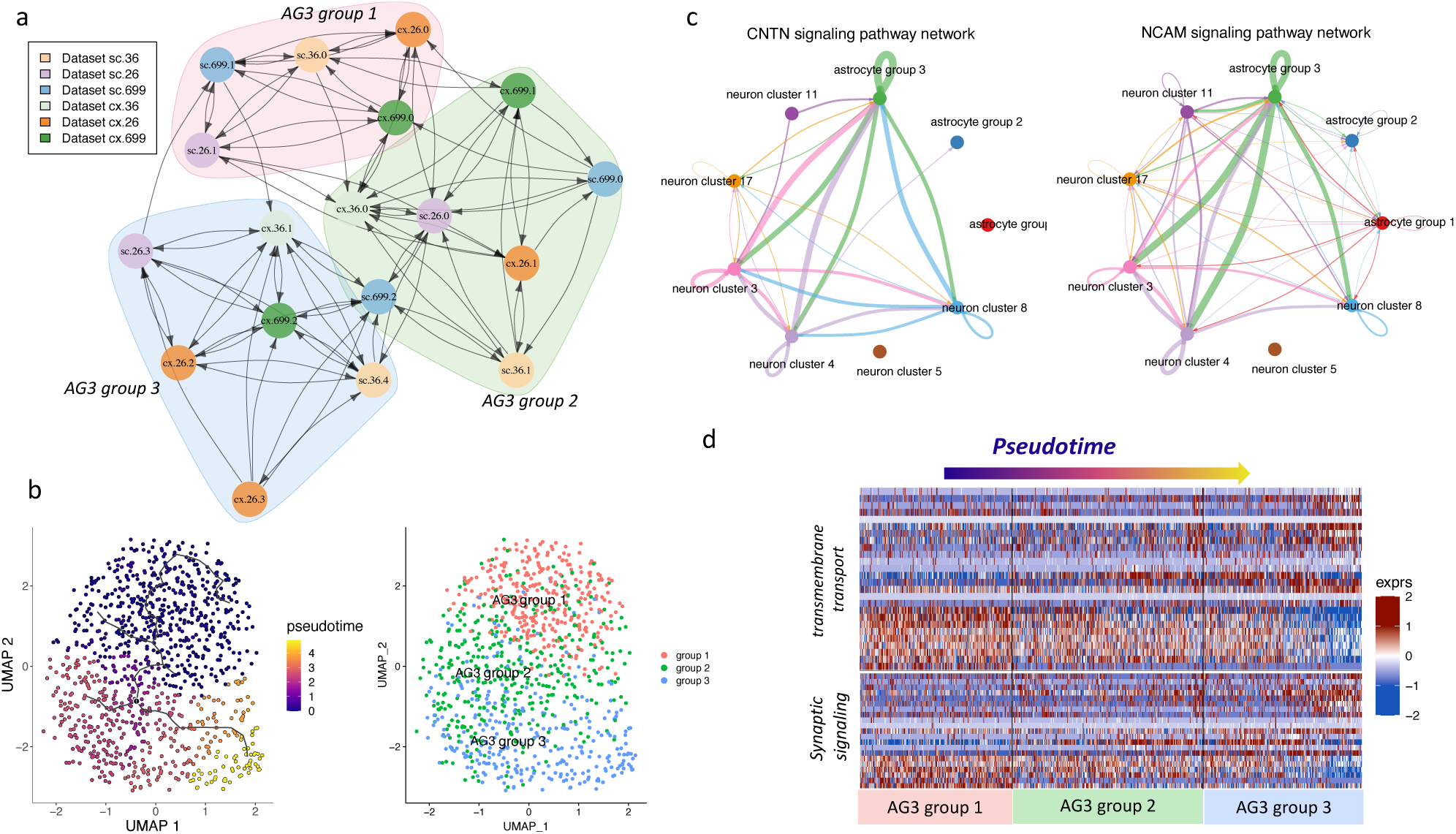
Detailed analysis of Astrocyte Group 3. **a.** Similarity of clusters within AG3 subpopulation. Community analysis over the graph showed three distinct groups. **b.** UMAP of AG3 subgroups and the computed pseudotime across these subgroups. Expression data from AG3 from all mice was previously merge and processed. **c.** Cell-cell signaling pathways CNTN and NEGR with an active role of AG3 with various neuron subclusters. **d.** Gene expression of relevant biological functions that showed significant differential expression as function of pseudotime.

## Discussion

Single cell sequencing represents a powerful approach offering an unprecedent resolution at molecular level. Here we introduce ClusterFoldSimilarity, a novel methodology designed to analyze collections of independent datasets including multimodal, multi-tissue or multi-species experiments, without data integration. Due to its interpretability, CFS enhance reproducibility and comparability of single cell studies, and can be easily integrated into the existing bioinformatics ecosystem. Our novel metric successfully identifies similar cell groups across any number of independent datasets, encompassing different sequencing technologies, achieving similar benchmarking scores at cell type labeling as other state of the art integrative methods, although with a higher accuracy when the number of overlapping features between single cell studies is low. Importantly, it achieves this without the need for an integrative step or batch effect removal over the sequencing signal, both of which can be strongly influenced by noise and inherent partial stochasticity in the data, potentially resulting in the loss of information. It first computes fold changes using a Bayesian approach to estimate differential abundance, adapting and expanding previous statistical frameworks, to construct a comparable frame of reference between groups and datasets. The similarity can be understood as a cumulative score across all features, in which concordant fold changes in the same vector space point at a similar phenotype. A potential limitation of our method arises when cell-level resolution is needed, as our metric operates at cell-group level, information at individual-cell level is not available. To partially mitigate this, incrementing the granularity and the number of groups could be desirable. Although, we show that our metric captures information related with the heterogeneity of cell groups, and the feature selection ability of our method can help identifying distinct markers of the composing members of the clusters on downstream analysis. We demonstrate the effective comparison of diverse omics datasets, first by benchmarking cell labeling accuracy using a complex mice gastrulation dataset, and a multi-species pancreatic assay. Secondly, we show how our methodology is capable of matching conserved cell groups across different species, using pancreatic single-cell RNA-seq from human and mouse. In a third example, we expand the multimodality to include single-cell proteomics (CyTOF), bulk-sorted RNA-Seq and single-cell RNA-Seq from synovial tissue of rheumatoid arthritis (RA) and osteoarthritis (OA) patients from B cells, T cells, monocytes, and stromal fibroblast populations. The similarity score correctly associated cell groups with the same phenotype, even with a small number of common features available (n=33). An additional multimodal assay was tested employing gene activity scores derived from ATAC-seq and a reference RNA-Seq data, matching populations across peripheral blood mononuclear cells (PBMC) datasets.

Finally, to exemplify a real use case, we sequence and analyze single-nuclei RNA-Seq data obtained from the spinal cord and motor cortex of adult mice, with a focus on exploring astrocytes as a disease-relevant glial cell subpopulation. By utilizing our similarity score, we identify three distinct subpopulations that are present in both tissues, each associated with diverse functionalities: a small population enriched in neurogenesis markers, a medium-size group of activated Gfap-positive astrocytes, and a large group of mature astrocytes with three additional sub-phenotypes. These findings will allow to further explore the roles of different astrocyte types or activation states depending, for example, on age or disease.

We believe that ClusterFoldSimilarity presents a valuable contribution to the ecosystem of bioinformatic tools for single-cell analysis, providing easily interpretable results, and solutions for the construction and exploitation of future cell atlases, which will be indispensable for gaining a comprehensive understanding of cell and tissue heterogeneity.

## Materials and methods

A typical single-cell data analysis workflow entails a series of essential steps. These steps commonly include filtering out low-quality cells and non-contributing features, normalizing and scaling the data, performing dimensionality reduction, and clustering or labeling cells based on groups defined by variability and phenotypic characteristics of interest.

Our method is designed to handle untransformed values (raw counts or peaks) and the specific groups of interest that need to be compared (e.g., clusters, cell types, cell states) in each independent dataset.

The features utilized for comparison can embody genes, transcripts, peaks, or other. Our workflow makes use of the set of features that are shared across all datasets. It is advisable to select features that exhibit some degree of variability or serve as markers of interest for our cell populations, albeit in some cases it might be desirable to include all available features when the phenotypic characteristics are unknown, or the main interest is that of marker selection.

### Fold change distribution and pairwise computation

Fold change can be formulated as the disparity in abundance between two sample sets or groups of samples/cells, and it is one of the most ubiquitous and intuitive metrics for evaluating hypotheses in quantitative experiments. Typically, we have a set of two samples representing a specific feature abundance, these can be seen as Poisson distributed random variables with mean μ, fold change is then formulated as the logarithmic ratio of the means of observed quantities such as mRNA fragments or protein levels, and statistical testing involves assessing the Null hypothesis that the measured entity exhibits no significant change in abundance under different conditions. Various statistical methods have been used and developed to calculate a P value associated with this measure of differential expression. However, caution must be exercised when interpreting measurements due to inherent technical biases and stochasticity of biological data. This concern is particularly pertinent when dealing with highly sparse single-cell data, where low or zero abundance observations may yield infinite or extremely high fold changes.

To mitigate this problem several approaches have been proposed, being the most widely used the addition of a small positive number p (often set as p=1) to the count data, known as a pseudocount. With this approach, problems like division by 0, or infinite *C_n_* values for log-normalization are avoided. But, assigning arbitrary numbers to p has a strong influence on the difference in abundance of low expressing entities: this is because artificially inflating the counts make disproportionate increase in low expressed features (for a feature with 1 count, adding a p=1 corresponds to a 100% increment, meanwhile a feature with expression 1000, a p=1 yields an increment in size of just a 0.1%). This has been demonstrated as problematic, both for differential expression and downstream analysis (81–84).

Here we extend the methodology devised by Erhard and Zimmer (84,85) which employs an empirical Bayes procedure to estimate pseudocounts and fold changes. We have made certain modifications to this approach (see details). We selected a fold change estimation procedure for several reasons: our similarity score is based on cumulative changes in abundances from a frame of reference (i.e. specific cluster or cell population), rather than the statistical significance of those changes. Consequently, we bypass the computation of statistical scores or P values. Furthermore, considering the large number of pairwise tests required to assess differential abundance and compute our similarity score, utilizing regression models to estimate size effects would incur in excessive computational time.

To calculate the pseudocounts for each group, we estimate the prior distribution based in the observed fold changes without pseudocounts by calculating:

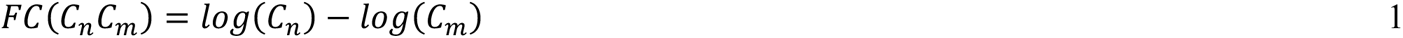

where *C_n_* and *C_m_* are respectively the mean of feature i expression in groups n and m. Consequently, we compute the mean *μ* and variance *σ*^2^of the prior as:

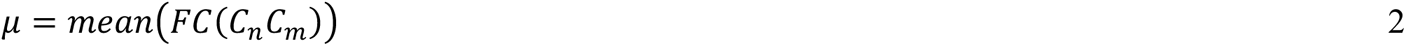

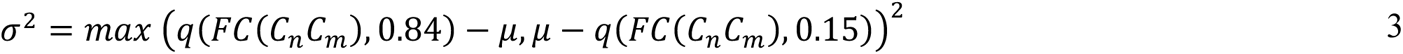

These quantiles correspond to one standard deviation for a Normal distribution. Importantly, to construct this prior distribution, we only use informative features, i.e. entries with non-zero values in both groups. Additionally, we impose a minimum of 3 features for calculating the prior, as opposed in the original (84) where 0 values are allowed on one group and no minimum number of informative features is required. If there are not sufficient values, we compute the FCs adding a pseudocount p (equation 4) to *C_n_* and *C_m_* for each gene, based on the hyperbolic arcsine (asinh) function with no log transformation, and we compute the variance using the standard formula. The asinh function has been shown to have desirable properties as pseudocount estimator (86,87), as it avoids preselecting an arbitrary constant, and yields a proportional pseudocount for each individual gene.

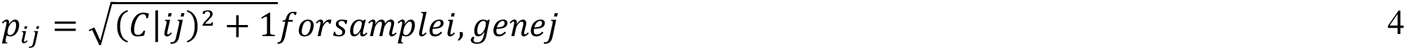

Finally, we select the mean instead of the median as the *μ* prior to reflect more extreme values in low abundance observations. In summary, we believe these changes are relevant when dealing with single-cell data, especially in the case of ATAC-Seq, single-nuclei RNA-Seq, and single-cell proteomics. As mentioned, *C_n_* and *C_m_* are expected to follow a Poisson distribution, being the conjugated prior of this distribution the gamma distribution. Following previous statistical frameworks (84), the expected mean and variance of the log distribution can be formulated as:

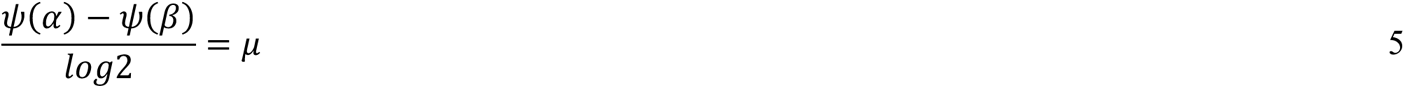

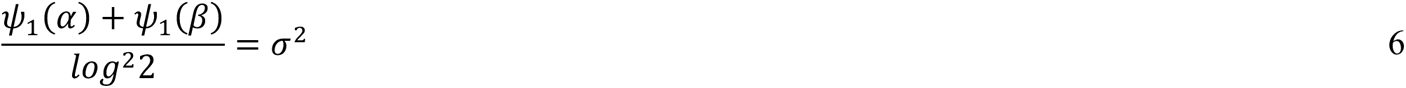

And thus, pseudocounts *α* and *β* can be computed by minimizing the function:

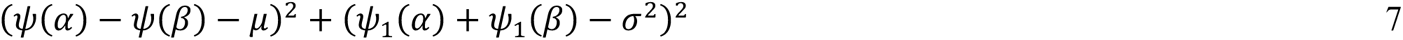

Where *ψ* and *ψ*_1_ are the digamma and trigamma functions respectively. Finally, the fold change distribution can be estimated using the new pseudocounts *α* and *β* for each feature as:

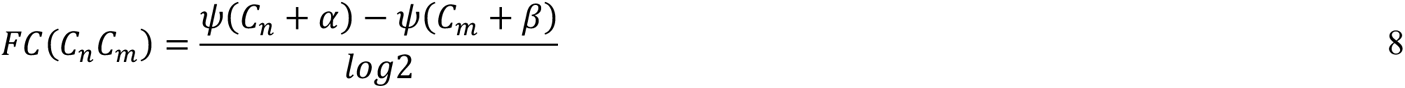

We apply the median to zero procedure (88) as a normalization factor over the new distribution of fold changes. This estimated FC is computed between cells in cluster/group i and cells in cluster/group j ∀*j* ≠ *i*. In such manner, we obtain k-1 FCs (k: number of clusters/groups in the dataset) for each feature (e.g., a dataset with 10 clusters/groups will yield 9 FC values for each feature on each of the 10 clusters/groups). By doing this we obtain a unique frame of reference for each cell group that encompass all the abundance differences from the perspective of each individual cell group, this information can be later compared with another external differential abundance reference, in our case, a cluster or group of cells coming from an independent dataset.

### Cell Subsampling and Fold Change Stabilization

As highlighted in preceding sections, a primary obstacle in calculating differential abundance arises from the occurrence of low to zero values, which can result in infinite or excessively high fold changes that do not accurately represent genuine biological effects. To mitigate this, we employ a permutation-based methodology to estimate the null distribution of the differences in abundance. Here, we randomly subsample 1/3 of the cells from each group “n” times and then estimate the fold changes utilizing the Bayesian approach (including the pseudocount estimation) described in the previous section:

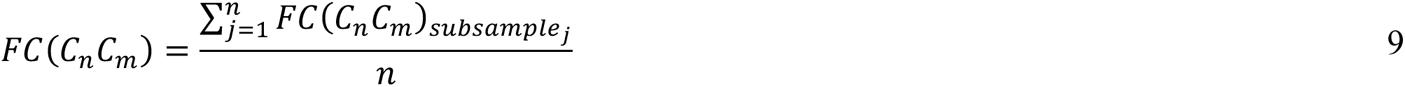

By subsampling 1/3 of the cells in each draw, the average number of permutations needed to observe all n cells in a group is:

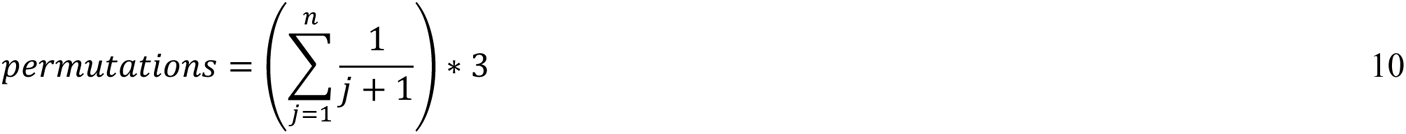

Which yields on average feasible computational times. As a reference, it is needed an average of 16 permutations to observe all cells in a group of 100 cells (Supplementary Fig. 3c).

Overall, we show (Supplementary Fig. 3a,d) that subsampling cells and estimating n times the fold change posterior distribution using a Bayesian procedure shrinks the mean of estimated FCs -coming from low count data and with no real and significant difference in abundance-towards zero, meanwhile maintaining the fold change distribution for significant values (Supplementary Fig. 3b,e). This is expected for a normal random distribution with mean *μ* = 0. By contrast, features with available expression data and true fold change maintain the same effect size throughout the n permutations.

*Supplementary Figure 3. Fold-change subsampling impact metrics using human pancreas alpha cells VS beta cells. **a** Comparison of non-significant fold-change values computed using Seurat and out methodology. **b** Comparison of significant (p.value.adj<0.01) fold-change values computed using Seurat and out methodology. For significant fold-changes the values are closer to the observed values by Seurat. c average number of subsamplings needed to observe every cell in a group of n. cells. d Distribution of non-significant fold-changes in the range [-0,2, 0.2] across the different subsampling sizes; 0 corresponds to the original observed log2 fold-change. e Distribution of significant (p.val.adj<0.01) fold-changes across the different subsampling sizes; 0 original observed log2 fold-change*.

### Similarity Score

We can think of fold-changes (FC) as a Euclidean vector with origin at 0, magnitude equal to the FC value, and direction equal to the sign of the expression change, the scalar product of two vectors in the same space gives as a result a perpendicular vector to the plane. In the case of the dot product of two FCs, the direction of the perpendicular vector will act as a concordance metric of the direction of the FCs (negative perpendicular vectors to the plane will indicate non-concordance in the differences in abundance of two frames of reference: the fold changes have opposite directions). The FCs are computed using the method described in the previous section, and it is calculated pairwise between each cluster i and all the rest of the clusters j belonging to the same dataset.

Once all frames of reference are computed (all the possible combinations of FCs between all clusters/groups from each independent dataset), we start by comparing the cluster 1 from dataset n with the cluster 1 in dataset m and continue iteratively. We achieve this comparison by computing, for each feature, the scalar product *S_i_* of every combination of FCs for the two clusters being compared:

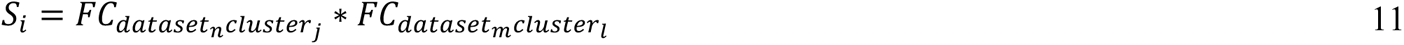

being the number of possible combinations *k*:

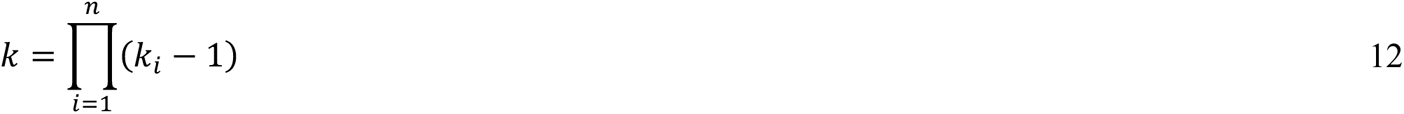

Being *k_i_*: the number of groups in dataset i. Then we calculate for each feature the mean of these k scalar products, that we denominate Scalar Contribution (SC):

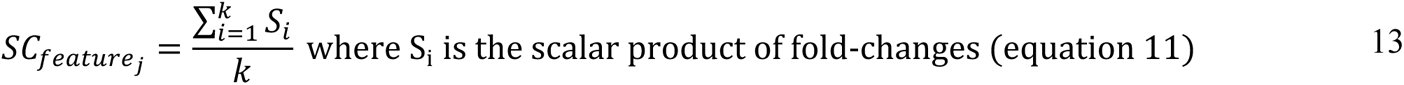

The assumption that we build upon is that two groups (e.g., clusters) resembling the same cell population will have distinct markers (either positive or negative), and thus FC for this marker must show similar size effects across independent datasets yielding a positive SC. This SC value will additionally serve as a measure of feature importance, being the feature with the highest positive value the one that contributes the most to the similarity between the two clusters (negative values of the scalar products indicate opposite FC values, and thus, dissimilarities between the changes in expression of the cell groups being compared). As these scores are built from differences in abundance, these selected contributing features can be seen as expression markers for both cell groups tied together by the similarity score.

An additional weight *ω* is added based on the concordance of the number of positive vs negative SC values (only values with absolute SC > 0.2 are considered), this assures to give more weight or penalize the similarity values considering the concordance of the direction of expression changes across datasets, especially for low FCs (a set of small negative SCs may add a small quantity to the overall similarity, but if the number of negative SCs is much larger than positive SCs this will indicate and overall tendency of non-concordant changes in abundances). Weight *ω* is calculated as:

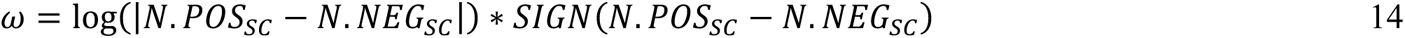

Finally, the similarity value for a pair of clusters is calculated as the square root of the sum (absolute value) of the Scalar Contributions from all features, retaining the sign of the sum:

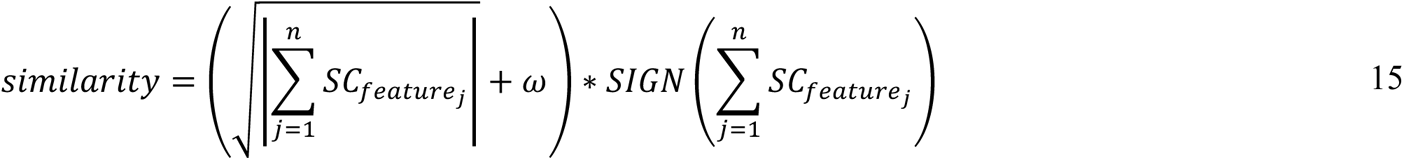

The highest similarity positive value calculated this way will correspond with the most similar cluster: it is expected that, for a specific population of cells, overall changes in gene expression across different clusters (other cell populations) are conserved on different datasets when using a similar frame of comparison, and thus the sign of the product of FCs should be positive and its module higher than non-concordant groups of cells.

### Similarity graph construction and community analysis

A directed graph is built from all the similarity results within the similarity table. Nodes represent cluster/groups defined by the user, and edges represent the similarity score. Edges are directed and point in the direction of the similarity score computed from cluster/group A to B meaning how similar is cluster A to cluster B, the weight (length) of edges correspond with the magnitude of the similarity score.

CFS can compute the graph with the n top similarities desired by the user, as an example, if only the top one similarity is obtained with CFS (option TopN=1), then each cluster n from dataset i (a node on the graph) will have only one outward edge (arrow) directed towards one cluster from dataset i+1 (another node of the graph). If the top two similarities are specified, each node will have two edges pointing outwards per external dataset. Once the graph has been built, we apply the Kamada-Kawai algorithm for graph visualization.

CFS can also perform a community analysis using the Leiden algorithm on the constructed graph, by using the package function *findCommunitiesSimmilarity* (which makes use of the igraph function graph_community_leiden), this analysis returns a table with the detected communities, and which clusters/groups correspond to each community, helping to aggregate the similar groups of clusters algorithmically, additionally the communities are depicted in different colors in the graph. The Leiden algorithm consists of three phases: (1) local moving of nodes, (2) refinement of the partition and (3) aggregation of the network based on the refined partition, using the non-refined partition to create an initial partition for the aggregate network.

### Mice motor cortex and spinal cord nuclei isolation and single nuclei RNA sequencing

Wild-type mice (C57BL/6NRj, Janvier Labs) were used in this study. Cortex was dissected from the left-brain hemisphere. Spinal cord was dissected and divided into three pieces. All tissues were flash frozen. For nuclei isolation, 20 µg of upper spinal cord (cervical / thoracic) and motor cortex were used. Samples were lysed in 2 ml of lysis buffer (5 mM CaCl_2_, 3 mM magnesium acetate, 0.32 M sucrose, 0.1 mM EDTA pH8.0, 10 mM Tris-HCl pH, 1 mM DTT, 0.5% Triton X-100) using a dounce homogenizer. Next, 3.7 ml of sucrose solution (1.8 M sucrose, 3 mM magnesium acetate, 1 mM DTT, 10 mM Tris-HCl pH) was pipetted to the bottom, and the ultracentrifugation tubes were filled up with the lysis buffer. Ultracentrifugation was performed at 24,400 rpm (107,164 rcf) for 2.5 hours at 4oC in a SW28 swing rotor (Beckman Coulter). Supernatant was discarded and the pelleted nuclei were resuspended on ice in 200 µl of DEPC-treated water-based 1X PBS. To get rid of any debris or nuclei aggregates, pre-separation filters of 30-µm pore size were used (Miltenyi Biotec). Nuclei were counted and the suspension was adjusted to 1,000 nuclei per microliter. For single-nuclei capture, 5,000 nuclei per sample were loaded on the Chromium controller (10x Genomics). All downstream steps were performed as described in the manufacturer’s protocol in the Chromium Next GEM Single Cell 3’ Kit v3.1. The libraries were sequenced on a NextSeq 550 System (Illumina) using the NextSeq 500/550 High Output Kit v2.5 (150 Cycles).

### Mice motor cortex and spinal cord single-nuclei RNA Seq bioinformatic analysis

Sequencing files from the 6 samples were processed using cellranger software version-6.0.1 and aligned to Genome Mouse Build 38 (mm10). Cells with more than 1.5% of mitochondrial content, library size inferior to 1500 counts, and less than 250 genes detected, were filter out. Downstream analysis was conducted individually using Seurat package on each of the six individual samples (tissue-mouse pairs). Cell type labeling was done using type specific markers from the scientific literature.

Astrocyte population was then subsetted for re-analysis, selecting clusters that showed unambiguous and distinct expression of astrocyte markers. The number of principal components for calculating nearest neighbors was chosen using both JackStraw and elbow methods in a data driven manner for each sample. Re-clustering was done using Louvaine algorithm, testing for granularity parameters that yield a coherent and stable number of clusters.

Similarity scores were obtained with ClusterFoldSimilarity statistical approach, using as a grouping variable the cluster id, and using a subsampling number of n=30. Feature selection of markers by CFS were obtained by specifying topN=10 (obtain top 10 markers contributing to the similarity for each cluster pair compared). Markers depicted in Figure 3c were selected as being a similarity contributor in more than one cluster pair contrast.

Differential expression between Astrocyte Groups (AG) was tested using a negative binomial model using the merged expression from all mice and tissues, and including as model covariates the tissue, the sequencing date, and the mouse id.

Cell-cell communication analysis was done using CellChat (89) version 2 (90) with default parameters, this method is based on literature-supported ligand-receptor interactions. Groups interrogated for cell-cell communication were obtained by clustering the neuronal populations using the merged data from all mice and only the motor cortex region, and the communication analysis was done between neuronal clusters and the three astrocyte populations described before.

Trajectory analysis, pseudotime scores, and graph analysis to identify differential expressed genes as a function of pseudotime for AG3 were obtained using Monocle2 (77) by using the raw merged expression of AG3 subpopulations from all mice and tissues and using Monocle2 default parameters. Gene set enrichment analysis was obtained with gprofiler2 R package.

### Benchmark of CFS and other integrative methods

Inspired by the approach used by Ghazanfar et. al. (10), we adapted extended and performed a benchmark comparison with other integrative and batch correction algorithms: MultiMap; naive PCA with no batch correction, PCA corrected using MNN (mutual nearest neighbors), and PCA corrected using Harmony; Seurat CCA (canonical correlation analysis); StabMap with no batch correction, StabMap corrected using MNN. We design two different scenarios: a scRNA-Seq pancreatic multispecies datasets (19), composed of mouse and human samples (human n=4, mice n=2), with different sizes of cell populations (human n=7561 cells, mice n=1861 cells), downloaded using R/Bioconductor package scRNAseq. The second scenario uses an assay of scRNA-seq that aims to study mouse gastrulation across entire embryos on E8.5 timepoint (20), downloaded using the MouseGastrulationData R/Bioconductor package (91) For each scenario, a pair of random samples were selected and processed using Seurat workflow (normalized, scaled, computation of top 2000 HVG, and dimensionality reduction), we then selected one sample as the reference dataset, to be used as ground-truth reference with the original annotation, and assign the other sample as the query dataset, keeping it unannotated and subjected to clustering analysis for the group similarity computation. Within each sampling round, we randombly selected a subgroup from the 2000 variable genes (n=25, 50, 100, 250, 500, 1000, 1500), doing this random sampling 4 times for each n gene subgroup (producing 4 observations of n genes between the same query and reference datasets). For StabMap, we followed the instructions depicted on the manual and benchmarking approach on the original paper, to compute the reference and query joint embedding into a common space, and using similar approaches with naive PCA, UINMF and MultiMAP.

To compute the accuracy at cell type label inference using CFS, clusters from the query dataset were labeled as the top-similar class group from the reference dataset, assuming that the entire cluster cell composition to be that cell type. Then accuracy was computed as assessing the number of cells from that cluster having the real annotation matching the assigned annotation. To assess the accuracy of the other integrative methods, we followed the approach by Ghazanfar et al., computing the mean accuracy of cell type annotation from the query sample cells using k-nearest neighbors with k = 5.

### Data Processing

#### Human and mouse single cell RNA-Seq from pancreas

Annotated data from both studies were downloaded using the R Bioconductor package scRNAseq. Downstream analysis was conducted using Seurat package on each of the datasets. The number of principal components for calculating nearest neighbors was chosen using both JackStraw (n=15). Clustering was done using Louvaine algorithm. ClusterFoldSimilarity was run using a subsampling of n=28. Barplots and exploratory analysis of the cell composition of clusters was done using LotOfCells R package (92). To analyze the correlation between similarity scores and cell type mixture composition of clusters, simulated single cell count data was produced using scDesign2 (93) selecting the Gaussian copula method to compute the model. The same number of cell per cell type were produced to account for population size (n=1200 cells per cell type).

#### RA and OR single-cell RNA-seq data, bulk RNA-seq data and mass cytometry data

Normalized and filtered data was downloaded from (https://www.immport.org/shared/study/SDY998. Expression was scaled, T-cell subpopulation was subsetted from the main dataset, re-analysed and re-clustered using Louvain algorithm (“FindNeighbors(reduction = “pca”, dims = 1:25)”, “FindClusters(resolution = 1)”). The same approach was followed for Cytof data with clustering resolution of 0.8.

PBMCs ATAC-Seq and single cell RNA-Seq:

We obtained two public single-cell datasets of human PBMCs. First, an unannotated scATAC-seq of human PBMCs were downloaded from the 10x Genomics website. In addition, a fully processed and high-quality annotated scRNA and ADT dataset was downloaded from https://atlas.fredhutch.org/data/nygc/multimodal/pbmc_multimodal.h5seurat (Hao et. al., 2021, PMID: 34062119) and used without further modifications. For the 10x Genomics dataset, the filtered peak-barcode matrices were loaded into R (v.4.0.0) using the Seurat (v.4.0.1)/Signac (v.1.4.0) functions “Read10X_h5” and “CreateChromatinAssay(min.cells = 10, min.features = 200. Barcodes from the unimodal dataset were filtered for a number of cutsites between 5,000 and 50,000 and a transcription start site score > 3.5. Gene activity scores were calculated using the function “GeneActivity” and log-normalized using as scaling factor the median of the gene activity matrix. Subsequently dimensionality reduction was applied based on the peak matrix using top 2000 variable, and cells were clustered using the first 20 PCA dimensions, with a resolution of 0.8. For the final cell type annotation, clusters were annotated based on marker gene expression. ClusterFoldSimilarity was run based on the gene activity matrix (10x Genomics dataset) or the RNA count matrix (ADT dataset) using all common features and fine cell type labels as cell identities, with a subsampling of n=30.

### Statistical significance

Throughout this study we considered a P-value threshold for statistical significance of 0.005, as it has been shown to foster reproducibility and lower the false positive rate (94,95). Therefore, any claims of significance on this paper should be noted as passing the threshold α = 0.005.

### Compatibility

ClusterFoldSimilarity is compatible with the most popular single-cell pipelines for an easy downstream integration: Seurat (96) and SingleCellExperiment (97).

## Availability of data and material

The algorithm ClusterFoldSimilariy is implemented in R language and freely available for download from Bioconductor repository (https://www.bioconductor.org/) as an R package (DOI: 10.18129/B9.bioc.ClusterFoldSimilarity), and on GitHub: https://github.com/OscarGVelasco/ClusterFoldSimilarity

Data from the pancreatic single-cell RNA-Seq datasets used for validation are available from Gene Expression Omnibus (GEO) under accession numbers: GSE84133 (98), GSE86469 (99), GSE81608 (100), ArrayExpress: E-MTAB-5061 (101)). PBMC ATAC-Seq and RNA-Seq data from 10x was downloaded from 10x genomic website: https://www.10xgenomics.com/resources/datasets/10k-human-pbmcs-atac-v2-chromium-controller-2-standard for unimodal ATAC-Seq. CITE-seq atlas of the circulating human immune system, comprising 160k cells with 26 annotated cell types was downloaded from GEO database under the accession number GEO: GSE164378. Rheumatoid arthritis (RA) and osteoarthritis (OA) single-cell RNA-seq data, bulk RNA-seq data and mass cytometry data were downloaded from ImmPort (https://www.immport.org/shared/study/SDY998) under study accession code SDY998.

## Acknowledgements

This publication was supported through state funds approved by the State Parliament of Baden-Württemberg for the Innovation Campus Health + Life Science Alliance Heidelberg Mannheim.

## Declaration of Interests

The authors declare no competing interests.

## Ethics approval

All animal experiments were performed in accordance with institutional guidelines of the University of Heidelberg and were approved by the local authority (Regierungspräsidium Karlsruhe, Germany).

## Notes

### Competing Interest Statement

The authors have declared no competing interest.

